# A fluorogenic complementation tool kit for interrogating lipid droplet-organelle interaction

**DOI:** 10.1101/2023.11.29.569289

**Authors:** Xiao Li, Rico Gamuyao, Ming-Lun Wu, Woo Jung Cho, Nathan B. Kurtz, Sharon V. King, R.A. Petersen, Daniel R. Stabley, Caleb Lindow, Leslie Climer, Abbas Shirinifard, Francesca Ferrara, Robert E. Throm, Camenzind G. Robinson, Alex Carisey, Alison G. Tebo, Chi-Lun Chang

## Abstract

Contact sites between lipid droplets and other organelles are essential for cellular lipid and energy homeostasis. Detection of these contact sites at nanometer scale over time in living cells is challenging. Here, we developed a tool kit for detecting contact sites based on <u>F</u>luorogen- <u>A</u>ctivated <u>B</u>imolecular complementation at <u>CON</u>tact sites, FABCON, using a reversible, low affinity split fluorescent protein, splitFAST. FABCON labels contact sites with minimal perturbation to organelle interaction. Via FABCON, we quantitatively demonstrated that endoplasmic reticulum (ER)- and mitochondria (mito)-lipid droplet contact sites are dynamic foci in distinct metabolic conditions, such as during lipid droplet biogenesis and consumption. An automated analysis pipeline further classified individual contact sites into distinct subgroups based on size, likely reflecting differential regulation and function. Moreover, FABCON is generalizable to visualize a repertoire of organelle contact sites including ER-mito. Altogether, FABCON reveals insights into the dynamic regulation of lipid droplet-organelle contact sites and generates new hypotheses for further mechanistical interrogation during metabolic switch.

## Introduction

Fatty acids are indispensable biomolecules to life as they are high-energy fuel for ATP production, building blocks for cell membranes, and signaling molecules for myriad biological functions. To harness such important molecules, eukaryotic cells develop an intricate inter-organelle network to dynamically stockpile and consume fatty acids upon demand. Lipid droplets are the center of this network and interact with many other organelles at contact sites to coordinate fatty acid metabolism (Henne et al., 2018; Olzmann and Carvalho, 2019; Walther et al., 2017). Contact sites are an evolutionarily conserved form of nanoscale spatial organization where two heterologous organelles are dynamically tethered to form close appositions with a gap distance of ∼20 nm (Gatta and Levine, 2017; Prinz et al., 2020; Scorrano et al., 2019). This nano-architecture exists between virtually all organelles to facilitate efficient and directed material transfer. Endoplasmic reticulum-lipid droplet (ER-LD) contact sites are essential for fatty acid storage in lipid droplets, while mitochondria-lipid droplet (mito-LD) contact sites are associated with fatty acid consumption and energy production upon nutrient deprivation (Renne and Hariri, 2021). Contact sites between peroxisomes and lipid droplets (PX-LD) have been shown to facilitate the elimination of lipid peroxides and maintain energy homeostasis during fasting (Binns et al., 2006; Chang et al., 2019; Kong et al., 2020). Overall, lipid droplet-organelle contact sites have diverse yet interrelated roles in fatty acid metabolism. Defects in lipid droplet-organelle interaction are associated with many metabolic disorders and neurological diseases, such as hereditary spastic paraplegia, ataxia, and early onset Parkinson’s disease (Herker et al., 2021).

Despite their importance and recent research endeavors, the dynamic regulation of lipid droplet-organelle contact sites remains poorly understood, primarily due to technical challenges in detecting these nanoscale foci. Light microscopy in conjunction with colocalization analysis between fluorescently labeled organelles is the most common method for indirect, collective readout when assessing organelle interaction within a cell. This LM-colocalization pipeline is easy to implement and allows the acquisition of statistically meaningful data across multiple temporal scales (Valm et al., 2017). Nonetheless, the spatial resolution of LM is insufficient for direct measurement of individual contact sites at nanoscale, further hindering the investigation of questions involved in how contact sites are dynamically distributed throughout a cell over time. In addition, the application of the LM-colocalization approach is often restricted to flat, adherent model cell lines, preventing our understanding of contact site biology in physiologically relevant cellular systems.

Assays based on enhanced signal upon proximity, such as bimolecular fluorescence complementation (BiFC) have been developed to detect contact sites via LM (Alford et al., 2012; Cieri et al., 2018; Eisenberg-Bord et al., 2016; Harmon et al., 2017; Kakimoto et al., 2018; Shai et al., 2018). The BiFC method requires protein-protein interaction of cognate split fluorescent proteins (FPs) localized on heterologous organelles at membrane juxtapositions and thus, faithfully report the location of contact sites. This design enables the BiFC method to be readily applicable to detect any organelle contact sites of interest. However, the implementation of BiFC assays often relies on traditional split FPs, which causes issues such as irreversible complementation and fluorescence leakiness without cognate partners. These issues may interfere with contact site dynamics and result in high background fluorescence, respectively (Bishop et al., 2019; Romei and Boxer, 2019; Tashiro et al., 2020).

We reasoned that a reversible BiFC reporter could eliminate the inherent issues associated with traditional split FPs and thus significantly improve its ability in detecting dynamic organelle interaction. A newly developed split FP system, splitFAST, appears to fit this purpose. This split system was engineered from the fluorescence-activating and absorption shifting tag (FAST), a 14-kDa apo-reporter that reversibly binds to hydroxybenzylidene rhodanine (HBR) analogs to become fluorescent (Plamont et al., 2016; Rakotoarison et al., 2023; Tebo and Gautier, 2019). HBR analogs are fluorogenic agents that strongly fluoresce when bound to splitFAST but are only weakly fluorescent in solution. The reversibility and high contrast signal from HBR analog binding make splitFAST an ideal reporter to implement BiFC for visualizing organelle contact sites with high spatial precision in living cells.

Here, we engineered and implemented the next-generation BiFC tool kit for quantitative visualization of lipid droplet-organelle contact sites using splitFAST (Figures 1A-B). We named this tool kit FABCON, for <u>F</u>luorogen-<u>A</u>ctivated <u>B</u>iFC at <u>CON</u>tact sites. To properly implement FABCON, we experimentally determined (i) the targeting mechanisms for localizing splitFAST to lipid droplets and other organelles and (ii) the affinity of splitFAST self-complementation to minimally affect organelle dynamics and proximity (Figure 1B). We first designed, generated, and validated a synthetic lipid droplet targeting motif based on Spastin’s lipid droplet targeting hairpin (Hp)(Chang et al., 2019). This synthetic Hp motif is highly enriched on lipid droplets and minimally affects lipid droplets’ functions. Next, we validated that splitFAST of low self-complementation (splitFAST_low_) (Rakotoarison et al., 2023) is suitable for implementing FABCON, as it did not affect organelle interaction. We confirmed that FABCON is completely reversible with no detectable fluorescence leakiness within living cells. Through the lens of FABCON, most lipid droplets are capable of making contact sites with the ER and mitochondria. Both ER-LD and mito-LD contact sites appear to be dynamic domains on lipid droplets’ surface and respond to distinct metabolic stimuli. While ER-LD contact sites were transiently increased during lipid droplet biogenesis, mito-LD contact sites were differentially and dynamically regulated during lipolysis and following inhibition of glycolysis inhibition. Via an automated line scanning algorithm, we quantitatively demonstrated distinct size of individual ER-LD and mito-LD contact sites. Furthermore, FABCON is generalizable to visualize intermittent PX-LD contact sites and frequent ER-mito interaction. Altogether, our data demonstrated that, with proper organelle targeting and affinity, FABCON is an effective tool kit that can be used to uncover the dynamic regulation of lipid droplet-organelle interaction and beyond.

**Figure 1.**
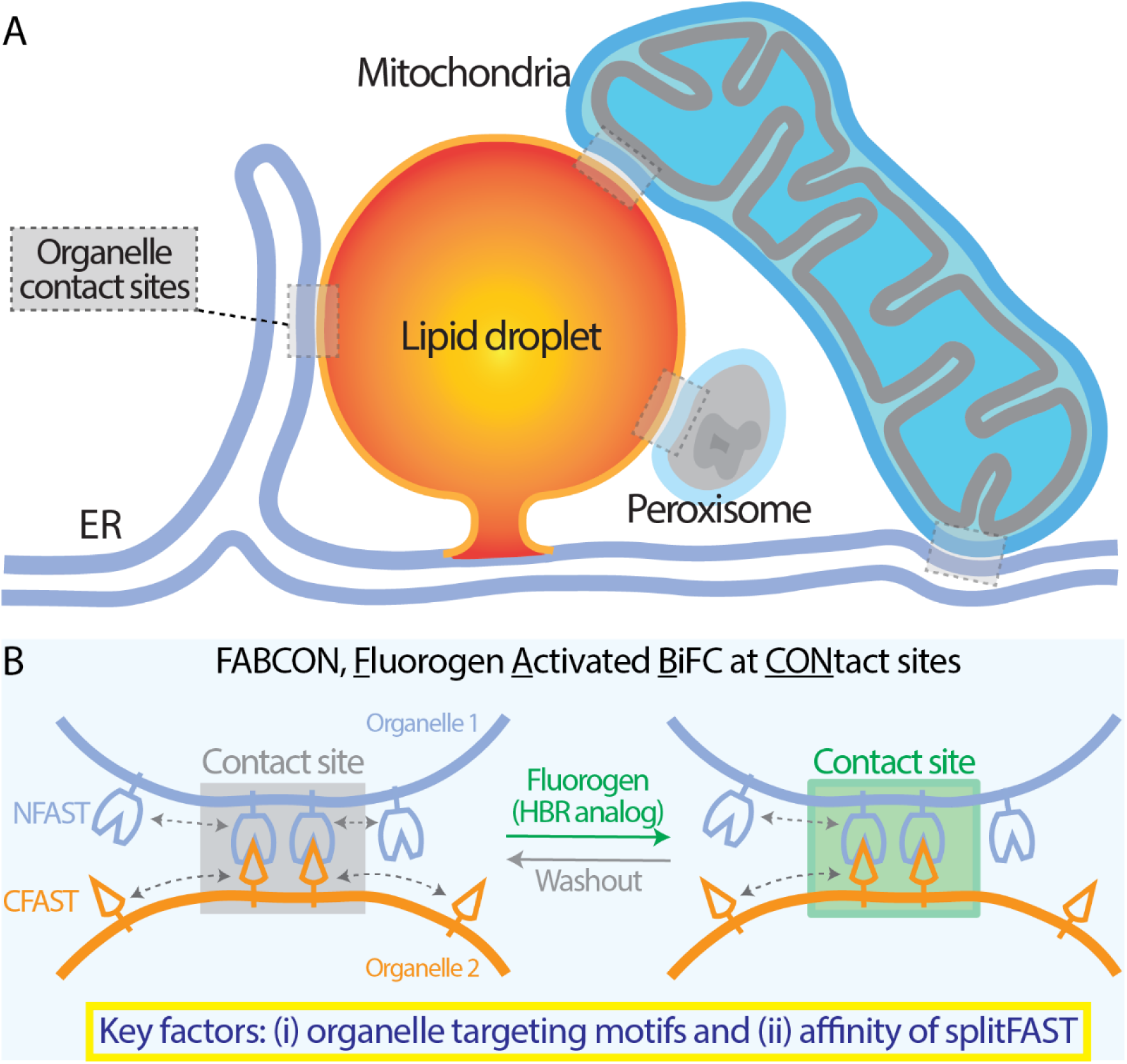
Visualization of lipid droplet-organelle contact sites using splitFAST-based FABCON. (A) Organelle contact sites between the endoplasmic reticulum and lipid droplets (ER-LD), mitochondria and LD (mito-LD), peroxisome and LD (PX-LD), and ER and mitochondria (ER-mito). (B) Diagram depicting the implementation of FABCON using a split reporter (splitFAST) and fluorogenic hydroxybenzylidene rhodamine (HBR) analog for detecting organelle contact sites.

## Results

### Engineering a synthetic lipid droplet targeting motif

Implementing FABCON for the detection of lipid droplet-organelle contact sites requires selective targeting splitFAST onto these organelles. Our previous study showed that Spastin’s Hp motif (1xHp, amino acid 43-92) has an affinity for lipid droplets (Chang et al., 2019). When overexpressed inside cells, this 1xHp motif was indiscriminately distributed between the ER and lipid droplets as detected by confocal microscopy in HeLa cells and by lipid droplet flotation assays of cell lysates from HepG2 hepatocyte (Figures 2A, 2B, and S1A). This dual organelle distribution of 1xHp motif hampered selective lipid droplet targeting. We reasoned that oligomerization of the Hp motif could increase its lipid droplet affinity (Chang et al., 2019), so we generated 6x tandem repeats of Hp (6xHp). In contrast to 1xHp, 6xHp was significantly enriched with lipid droplets with negligible ER localization (Figures 2A-C). The higher affinity of 6xHp was further demonstrated by fluorescence recovery after photobleaching (FRAP) analysis on lipid droplets in U2OS cells. A moderate recovery was detected for the fluorescence of 1xHp because it can diffuse between the ER and lipid droplets. In contrast, only minimal recovery for the 6xHp fluorescence was observed following photobleaching (Figures 2D-F), suggesting 6xHp was primarily static on lipid droplets’ membrane. Furthermore, structured illumination microscopy (Scorrano et al.) with twofold increase in resolution revealed that 6xHp localized to lipid droplets rather than to nearby ER membranes labeled with mEmerald-Sec61β (Figure 2G), supporting the idea that 6xHp directly localizes to the lipid droplet membrane monolayer. Consistent with this, we observed strong electron-dense precipitates juxtaposed to lipid droplets via electron microscopy (EM) in APEX2-6xHp overexpressing U2OS cells following H_2_O_2_ treatment (Figure 2H). Altogether, these results indicate that 6xHp is highly enriched on lipid droplets in multiple cell types and thus, making it a suitable targeting motif to bring reporters onto these organelles.

**Figure 2.**
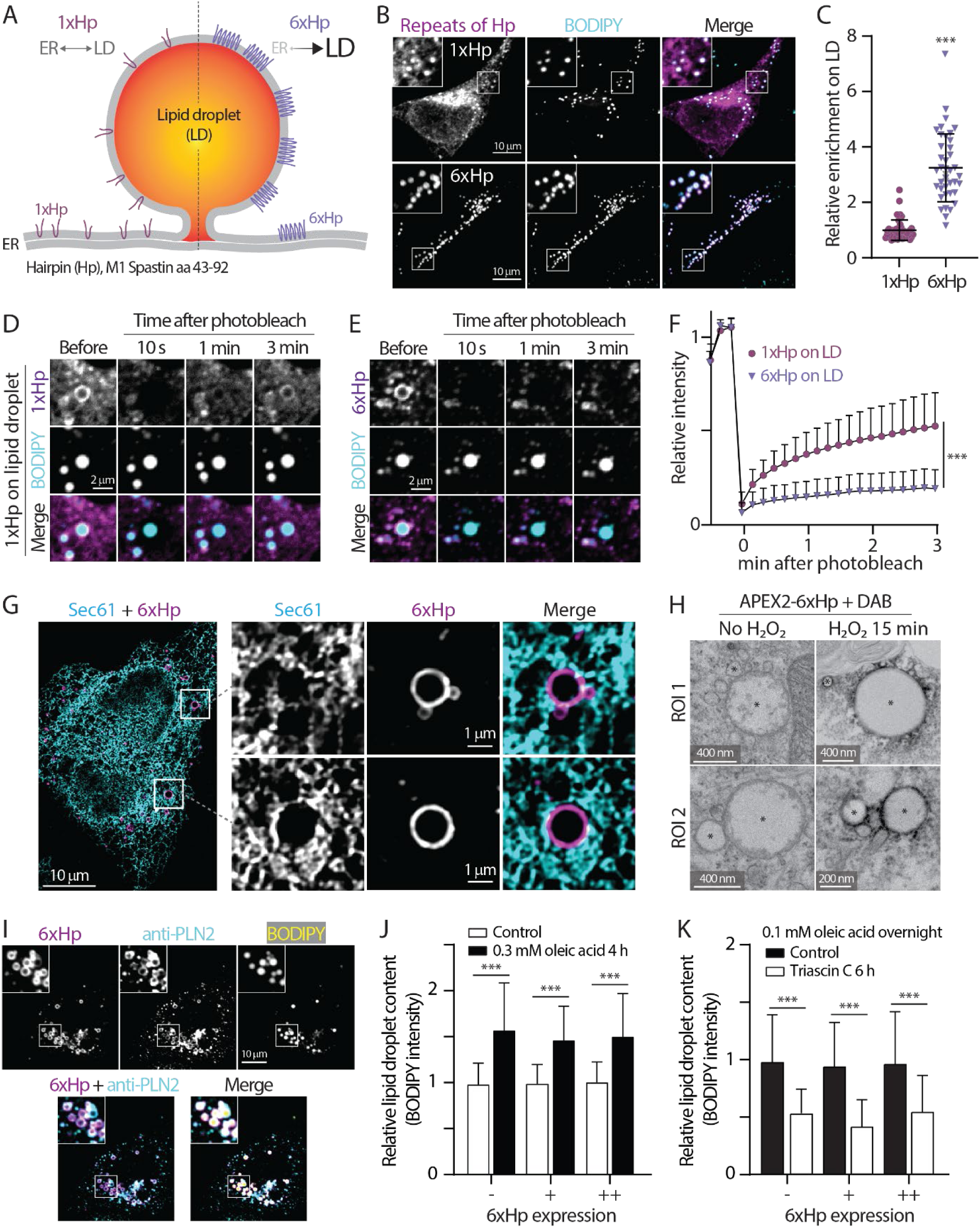
The design and validation of a synthetic lipid droplet targeting motif. (A) Diagram depicting the ER and lipid droplet (LD) distribution of 1xHp (hairpin; amino acids 43-92 from human M1 Spastin) and 6xHp. (B) Localization of BODIPY 493/503-labeled LDs and mApple-1xHp or mApple-6xHp in HeLa cells treated with 100 µM oleic acid (OA) overnight. Maximal intensity projected (MIP) confocal images from six axial slices (1.8 µm in total thickness) are shown. (C) Quantification of relative enrichment of 1xHp and 6xHp on LD from (B). Raw data and mean ± standard deviation are shown (53-56 cells from three independent experiments). (D and E) Fluorescence recovery after photobleaching (FRAP) of 1xHp (D) and 6xHp (E) on LDs in OA-treated U2OS cells labeled with BODIPY monitored by confocal microscopy. (F) Quantification of FRAP of (D and E). Mean ± standard deviation are shown (22-37 cells from 3-4 independent experiments). (G) Subcellular localization of Halo-6xHp relative to ER marker mEmerald-Sec61β in an OA-treated HeLa cell monitored by structured illumination microscopy. MIP images from ten axial slices (∼2 µm in total thickness) are shown. (H) Electron micrographs of LDs in U2OS cells expressing APEX2-6xHp incubated with DAB in the absence or presence of H_2_O_2_. * indicates representative LDs. (I) Colocalization of Halo-6xHp and endogenous LD protein PLIN 2 in an OA-treated U2OS cell stained with BODIPY monitored by confocal microscopy. Representative images from a single axial plan (0.3 µm) are shown. (J) Relative LD content in Halo-6xHp overexpressing HeLa cells under control condition or treated with 0.3 mM OA for 4 h. Mean ± standard deviation are shown (98-217 cells from three independent experiments). ‘-‘ indicates absence of 6xHp expression. + and ++ indicate low and moderate expression of 6xHp, respectively. (K) Relative LD content in OA-loaded, Halo-6xHp overexpressing HeLa cells under control condition or incubated with 10 µM Triacsin C for 6 h. Mean ± standard deviation are shown (169-499 cells from three independent experiments). ‘-‘ indicates absence of 6xHp expression. + and ++ indicate low and moderate expression of 6xHp, respectively.

Overexpressing synthetic 6xHp may affect native proteins’ ability to access lipid droplets via protein crowding (Kory et al., 2015) and compromise functions of these organelles. We first eliminated these concerns by demonstrating that an endogenous lipid droplet protein, perilipin 2 (PLIN 2) remained on lipid droplets decorated with overexpressed 6xHp (Figure 2I). In addition, overexpressing 6xHp at low and moderate levels (similar levels for all imaging experiments) minimally affected lipid droplet biogenesis (Figure 2J), lipid droplet breakdown following inhibition of long chain acyl-CoA synthase by Triacsin C treatment (Figure 2K)(Hartman et al., 1989; Roberts et al., 2023; Tomoda et al., 1987), and lipid droplet consumption incubation with Hanks’ balanced salt solution (Figure S1B). In conclusion, we found that 6xHp is suitable for targeting lipid droplets with negligible perturbation to the functions of these organelles. Targeting motifs for other organelles, including the ER, mitochondria, and peroxisomes, were also validated via colocalization analysis with known organelle markers (Table S1).

### Experimental determination of low affinity splitFAST for implementing FABCON

Interaction of splitFAST from heterologous organelles constitutes a tether for contact sites. We reasoned that the affinity between the cognate pair of splitFAST is crucial to implementing FABCON for detecting, but not creating, organelle interaction. All experiments for testing the effects of splitFAST with distinct affinity on organelle interaction were performed in the absence of fluorogen. We first examined whether high affinity splitFAST (the original splitFAST) with a self-complementation *K*_d_ of ∼3 µM (Rakotoarison et al., 2023; Tebo and Gautier, 2019) drives organelle interaction within cells. We tagged high-affinity NFAST to the N-terminal of mApple-6xHp (NFAST_high_-LD) and CFAST to the N-terminal of Halo-ER (CFAST-ER) to detect the effect of NFAST_high_-CFAST interaction on these organelles. We found that a subpopulation of CFAST-ER was enriched with NFAST_high_-LD, suggesting enhanced ER-lipid droplet interaction (Figure 3A). FRAP analysis further revealed that recovery of CFAST-ER in regions near NFAST_high_-LD was significantly attenuated than those in the bulk of ER, indicating that CFAST-ER was indeed trapped at ER-LD contact sites via NFAST_high_-LD interaction (Figure 3B). In contrast, low affinity splitFAST with a self-complementation *K*_d_ of 220 µM (Rakotoarison et al., 2023) did not lead to CFAST-ER colocalizing with NFAST_low_-LD (Figure 3C). Moreover, CFAST-ER FRAP in regions near NFAST_low_-LD was similar to in those in the ER (Figure 3D). These results indicated that NFAST_high_-CFAST may lead to the expansion of contact sites while NFAST_low_-CFAST only marginally affects organelle interaction. Consistent with this finding, NFAST_high_-peroxisome (PX-NFAST_high_) displayed significantly higher colocalization with CFAST-LD while PX-NFAST_low_ and CFAST-LD were sporadically distributed within cells and make intermittent interactions as in control condition (Figures S2A-B). Similar organelle interaction patterns were also observed between lipid droplet and mitochondria: NFAST_high_-CFAST significantly enhanced interaction as compared to NFAST_high_ alone or NFAST_low_-CFAST (Figures 3E and 3F). Importantly, these results demonstrated that, when confined to a membrane (a 2-dimensional surface), the reversible bimolecular interaction with sub-µM *K*_d_ can drastically interfere with organelle interaction and distribution. Direct measurement of the length of mito-LD contact sites via EM further confirmed that NFAST_low_-CFAST had no effect on the size of contact sites (Figure 3G). Based on the analyses above, we decided to implement FABCON with the NFAST_low_-CFAST pair, as it minimally affects organelles’ interaction.

**Figure 3.**
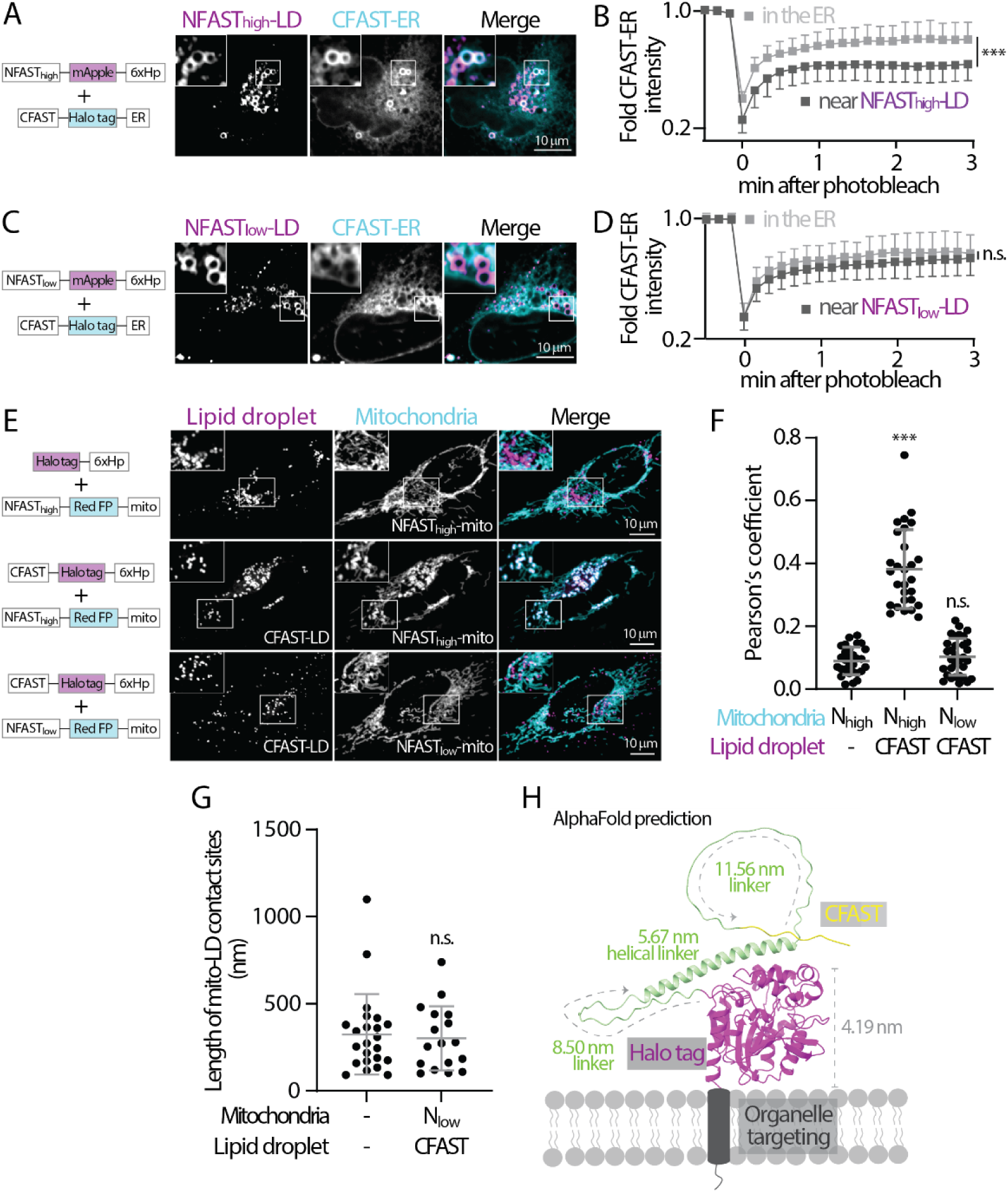
Low affinity splitFAST is suitable for implementing FABCON. (A) Distribution of NFAST_high_-mApple-6xHp and CFAST-Halo-ER in an oleic acid (OA)-treated U2OS cell. Representative images from a single axial plan are shown. (B) Quantification of FRAP of CFAST-Halo-ER in the ER and near NFAST_high_-decorated lipid droplets. Mean ± standard deviation are shown (24-25 regions from three independent experiments). (C) Distribution of NFAST_low_-mApple-6xHp and CFAST-Halo-ER in an OA-treated U2OS cell. Representative images from a single axial plan are shown. (D) Quantification of fluorescence recovery after photobleaching (FRAP) of CFAST-Halo-ER in the ER and near NFAST_low_-decorated lipid droplets. Mean ± standard deviation are shown (20-22 regions from three independent experiments). (E) Distribution of lipid droplet and mitochondria in OA-treated HeLa cells overexpressing Halo-6xHp and NFAST_high_-mApple-mito (top), CFAST-Halo-6xHp and NFAST_high_-mApple-mito (middle), or CFAST-Halo-6xHp and NFAST_low_-mApple-mito (bottom) monitored by confocal microscopy. Maximal intensity projected images from six axial slices (1.8 µm in total thickness) are shown. (F) Quantification of the Pearson’s colocalization coefficient of lipid droplet and mitochondria described in (E). Raw data and mean ± standard deviation are shown (29-32 cells from three independent experiments). (G) Quantification of the length of mito-LD contact sites detected by SEM in control and in FAB^mito-LD^ (see Figure S4B) infected HeLa cells. Raw data and mean ± standard deviation are shown (n=23 in control; n=17 in N_low_+CFAST). (H) AlphaFold structure prediction of CFAST-linker-Halo displayed on a membrane bilayer. The size of Halo tag and length of flexible and helical linkers are indicated.

We further confirmed that there is no fluorescence leakiness in this system. The fluorogen signal was only observed when both CFAST-LD and NFAST_low_-ER were present within cells, but not in cells with CFAST-LD or NFAST_low_-ER alone (Figures S3A-B). The appearance of this fluorogen signal complemented by NFAST_low_-CFAST occurred rapidly following the addition of the fluorogen and promptly disappeared after the washout (Figure S3C), indicating that the splitFAST interaction at contact sites is completely reversible. In conclusion, our data demonstrated that NFAST_low_-CFAST is ideal for implementing FABCON to detect dynamics of contact sites in living cells.

In addition, we inserted flexible and helical linkers to the CFAST-containing halves ensuring proper spatial accommodation for splitFAST interaction from heterologous organelles. Based on AlphaFold structure prediction (Jumper et al., 2021), CFAST could explore the space of 5 nm to 30 nm from the organelle membrane when the linkers are fully extended (Figure 3H). To better control the expression levels of the cognate pairs of splitFAST, we cloned them into a bicistronic IRES backbone and generated lentiviral particles of these IRES plasmids for cellular delivery (Figure S4 and Table 1). Moreover, we kept the Halo tag in the CFAST-containing halves for analysis purposes but replaced the FP in the NFAST-containing halves with spacers and an epitope tag to allow multiplex imaging. The expression and localization of all FABCON pairs were validated via LM (Figures S4A-D). For visualization and quantification of contact sites, we applied only 3 µM or lower of HBR-2,5DOM (Mineev et al., 2021) as fluorogen to circumvent issues of enhancing contact sites formation associated with higher concentration of fluorogen.

**Table 1.** SplitFAST pairs for contact sites.

| Contact sites | NFAST half | CFAST half | IRES plasmid (5'-3') |
| --- | --- | --- | --- |
| ER-LD | NFAST<sub>low</sub>-ER | CFAST-LD | NFAST<sub>low</sub>-ER_IRES_CFAST-L-LD |
| Mito-LD | NFAST<sub>low</sub>-mito | CFAST-LD | NFAST<sub>low</sub>-miito_IRES_CFAST-L-LD |
| PX-LD | PX-NFAST<sub>low</sub> | CFAST-LD | PX-NFAST<sub>low</sub>_IRES_CFAST-L-LD |
| ER-mito | NFAST<sub>low</sub>-ER | CFAST10-mito | NFAST<sub>low</sub>-ER_IRES_CFAST10-L-mito |

ER: endoplasmic reticulum; LD: lipid droplet; mito: mitochondria; PX: peroxisome; L: linker

### FABCON reveals the dynamic regulation of ER-LD contact sites

We first examined ER-LD contact sites using FABCON (FAB^ER-LD^; see Figure S4A). In live HeLa cells, ER-LD contact sites appeared to be prevalent and were present on 93% of lipid droplets labeled by CFAST-LD (Table 2), suggesting most lipid droplets are able to form contact sites with the ER. Some lipid droplets had full coverage of ER-LD contact sites while others displayed distinguishable domains and are only partially covered by these contact sites (Figure 4A), indicating heterogenous distribution of ER-LD contact sites within the lipid droplet population. These properties were consistent in U2OS cells, where 96% of lipid droplets were able to form ER-LD contact sites; and 46% among those were partially covered by contact sites (Table 2). These contact site domains can be dynamic as they appear to move along the lipid droplet’s surface and change their distribution and intensity within minutes (Figure 4B). Lattice light sheet microscopy (LLSM) provided additional insights into these domains in 3D (Figure 4C): whereas lipid droplets’ surface is smooth and continuous, ER-LD contact sites displayed uneven distribution and distinct domain structures. Notably, the 3D structure of ER-LD contact sites on each lipid droplet differed from others. As confocal microscopy does not have sufficient spatial resolution, full coverage suggests (i) ER completely wraps around lipid droplet’s perimeter or (ii) individual ER-LD contact sites that were not visualized with confocal microscopy.

**Figure 4.**
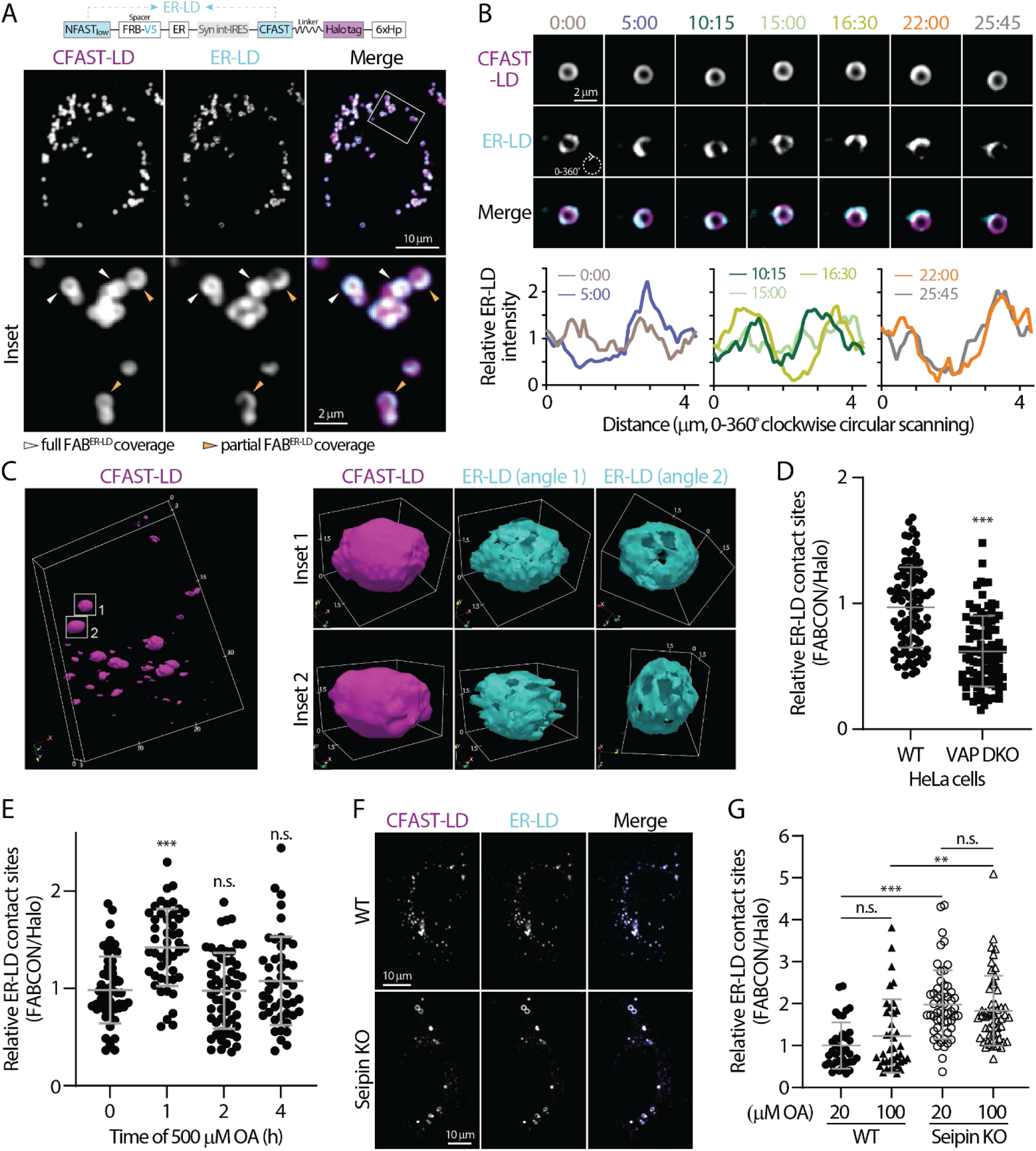
Dynamic regulation of ER-LD contact sites revealed via FAB^ER-LD^. (A) Detection of lipid droplets and ER-LD contact sites in OA-treated HeLa cells producing FAB^ER-LD^ (top) monitored by confocal microscopy. Representative maximal intensity projected images from three axial slices (∼1 µm in total thickness) are shown. (B) Dynamics of ER-LD contact sites on a lipid droplet in OA-treated U2OS cell producing FAB^ER-LD^ monitored by confocal microscopy over time (top). Relative intensity profiles of ER-LD measured by clockwise circular scanning are shown in bottom panels. (C) 3D rendering of lipid droplets and ER-LD contact site in U2OS cells imaged via LLSM. Lipid droplets labeled by CFAST-LD from a whole cell are shown on the left. Box volume is 24.6 μm x 16.8 μm x 2.67 μm. Two individual lipid droplet and their ER-LD contact sites are shown on the right. Box volumes are 3.10 μm x 2.35 μm x 2.52 μm (top row) and 3.28 μm x 2.40 μm x 2.82 μm (bottom row). Numbers indicate μm. (D) Relative levels of ER-LD contact sites in control and VAP DKO HeLa cells. Raw data and mean ± standard deviation are shown (96 cells for each condition from three independent experiments). (E) Relative levels of ER-LD contact sites HeLa cells pulsed with 500 μM of OA over time. Raw data and mean ± standard deviation are shown (47-58 cells from three independent experiments). (F) ER-LD contact sites in oleic acid (OA)-treated WT and Seipin KO SUM159 cells producing FAB^ER-LD^ monitored by confocal microscopy. Representative maximal intensity projected images from three axial slices (∼1 µm in total thickness) are shown. (G) Relative levels of ER-LD contact sites in WT and Seipin KO SUM159 cells producing FAB^ER-LD^ and treated with 20 or 100 µM OA overnight. Raw data and mean ± standard deviation are shown (37-55 cells from four independent experiments).

**Table 2.**
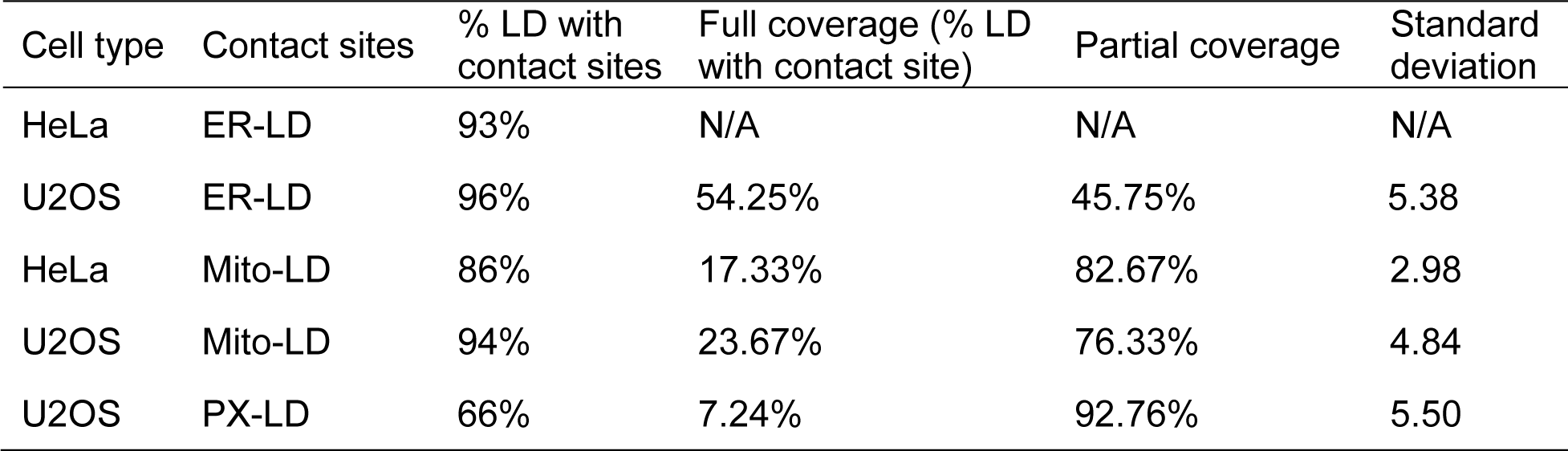
Contact sites statistics.

| Cell type | Contact sites | % LD with contact sites | Full coverage (% LD with contact site) | Partial coverage | Standard deviation |
| --- | --- | --- | --- | --- | --- |
| HeLa | ER-LD | 93% | N/A | N/A | N/A |
| U2OS | ER-LD | 96% | 54.25% | 45.75% | 5.38 |
| HeLa | Mito-LD | 86% | 17.33% | 82.67% | 2.98 |
| U2OS | Mito-LD | 94% | 23.67% | 76.33% | 4.84 |
| U2OS | PX-LD | 66% | 7.24% | 92.76% | 5.50 |

ER-LD contact sites are tethered via many protein complexes, including VAP-VPS13 interaction (Kumar et al., 2018; Murphy and Levine, 2016). Thus, we validated the level of ER-LD contact sites via FAB^ER-LD^ in VAP-A and VAP-B double knockout (VAP DKO) HeLa cells (Figure S5A). We used the ratio of the FAB^ER-LD^ signal to Halo signal (expression level normalization) as a relative readout for ER-LD contact sites and observed a significant decrease in ER-LD contact sites in VAP DKO cells as compared to parental HeLa cells (Figure 4D). This result indicated that FAB^ER-LD^ is suitable for detecting ER-LD contact sites at an endogenous level because it faithfully reports the expected changes in VAP DKO cells.

We next investigated how ER-LD contact sites are regulated during lipid droplet biogenesis in HeLa cells. Following 500 µM oleic acid (OA) acid incubation, we observed a significant (45%) increase in ER-LD contact sites at 1 h (Figure 4E), reflecting the correlation of these sites in lipid droplet formation (de Vries et al., 1997; Fujimoto et al., 2007). Intriguingly, this increase was transient as ER-LD contact sites decreased and returned to the baseline level after 2h (Figure 4E). These results indicated that ER-LD contact sites are dynamically regulated during lipid droplet biogenesis.

Seipin is an ER membrane protein regulating lipid droplet size and distribution during biogenesis via bridging the membrane continuum between the ER and lipid droplets (Fei et al., 2011; Fei et al., 2008; Salo et al., 2016b; Salo et al., 2019; Wang et al., 2016). We wondered if seipin plays a role in the formation of ER-LD contact sites. Consistent with previous studies (Salo et al., 2016a; Wang et al., 2016), we found that lipid droplets in seipin knockout (KO) were heterogenous is size with a few larger ones compared to wild-type cells (Figures 4F and S5B). Intriguingly, we observed a significant increase in ER-LD contact sites in seipin KO cells either under low (20 µM) or high (100 µM) OA incubation that mimic resting state and fatty acid surplus conditions, respectively. Altogether, FABCON was able to detect ER-LD contact sites with high spatial precision and revealed dynamic changes in these sites following a variety of stimuli and manipulations.

### Dynamic regulation of mito-LD contact sites during metabolic switching

We next characterized mito-LD contact sites using FAB^Mito-LD^ (Figure 5A). Consistent with ER-LD contact sites, ∼90% lipid droplets were competent to make contact sites with mitochondria in HeLa and U2OS cells (Table 2). Interestingly, among these lipid droplets able to form contact sites, only ∼20% were fully covered with mito-LD contact sites while the other 80% often displayed distinct contact site domains (partial coverage). These observations demonstrated that mito-LD contact sites were less prevalent on a lipid droplet surface compared to ER-LD contact sites. Moreover, mito-LD contact sites often outlined the perimeter of lipid droplet clusters but are less common in the center region of these clusters (Figure 5A). Some mito-LD contact site domains were dynamic as they appeared to fuse and move along the lipid droplet’s surface within minutes (Figure 5B).

**Figure 5.**
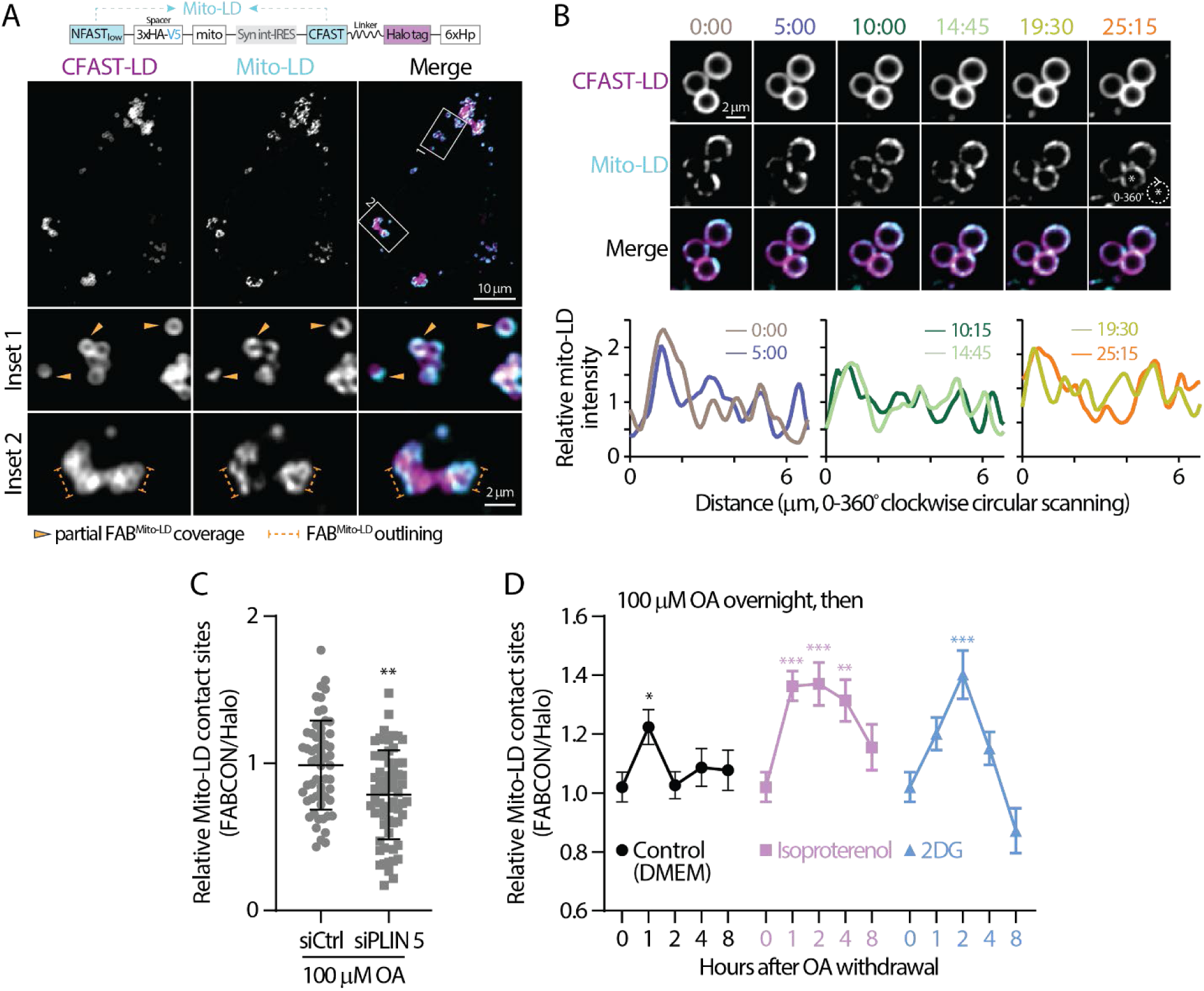
Dynamic regulation of mito-LD contact sites revealed via FAB^mito-LD^. (A) Detection of lipid droplets and mito-LD contact sites in oleic (OA)-treated HeLa cells producing FAB^mito-LD^ (top) monitored by confocal microscopy. Representative maximal intensity projected images from two axial slices (0.6 µm in total thickness) are shown. (B) Dynamics of mito-LD contact sites on a lipid droplet in OA-treated U2OS cell producing FAB^mito-LD^ monitored by confocal microscopy over time (top). Relative intensity profiles of mito-LD measured by clockwise circular scanning are shown in bottom panels. (C) Relative levels of mito-LD contact sites in OA-treated HeLa cells transfected with scramble or PLIN 5 siRNA. Raw data and mean ± standard deviation are shown (61-63 cells from three independent experiments). (D) The temporal dynamics of mito-LD contact sites in OA-treated HeLa cells following OA withdrawal in DMEM and in DMEM with 10 µM of isoproterenol or 4 mM 2DG. Mean ± standard error are shown (37-58 cells from 3-4 independent experiments). Statistical significance was compared to time zero.

Recent studies demonstrated that Perilipin 5 (PLIN 5) is a tether for mito-LD contact sites (Miner et al., 2023b; Ouyang et al., 2023; Wang et al., 2011). Therefore, we examined the levels of mito-LD contact sites in PLIN 5 knockdown HeLa cells (Figure S5C). We found a moderate yet significant reduction in mito-LD contact sites in siPLIN 5-treated HeLa cells following 100 µM OA incubation (Figure 5C), suggesting that PLIN 5 tethers lipid droplets to mitochondria under fatty acid surplus conditions. In contrast, wild-type and VAP DKO HeLa cells showed comparable levels of mito-LD contact sites (Figure S6A), as VAP-A and VAP-B are ER-resident proteins that have no direct role in mito-LD tethering. In addition, knocking out an PX-LD tether, spastin, had no effect on mito-LD contact sites (Figures S5D and S6B). These data suggest that mito-LD contact sites are independently maintained regardless of the status of other lipid droplet-organelle interaction.

Mitochondria are essential for fatty acid oxidation (FAO) to generate ATP. We reasoned that mito-LD contact sites may be regulated when switching to fatty acid oxidation (FAO)-dependent conditions as these sites can facilitate fatty acid trafficking. To test this idea, we measured mito-LD contact sites following OA withdrawal in HeLa cells treated with 100 µM OA overnight. We observed a transient increase in mito-LD contact sites 1 h after incubating these OA-loaded cells control media (Figure 5D). In contrast, a sustained increase in mito-LD contact sites was observed in cells treated with 10μM isoproterenol (Figure 5D), a known drug to induce lipolysis (Gallardo-Montejano et al., 2016). These results indicated that the extent of mito-LD contact sites was correlated with fatty acid release from lipid droplets. Interestingly, treatment with 2DG, which inhibits glycolysis and likely enhances FAO (Brown, 1962; Shiratori et al., 2019; Sottnik et al., 2011), during OA withdrawal ultimately resulted in a peaked increase in mito-LD contact sites at 2 h and eventually dropped below baseline after 8 h (Figure 5D). Altogether, these data revealed differential regulation of mito-LD contact sites under various metabolic conditions.

### Automated contact site analysis

The domains of ER-LD and mito-LD contact sites could be easily observed on larger lipid droplets in U2OS cells. To further quantify these domains, we developed an automated line-scanning analysis pipeline for unbiased <u>co</u>ntact <u>si</u>te <u>ma</u>pping (COSIMA). The COSIMA pipeline involves: (i) a machine learning-based identification of the outline and void of lipid droplets using Ilastik, (ii) a Python-based traverse algorithm to measure the intensity of FABCON (contact site) and Halo (lipid droplet) channels along the edges of the voids, and (iii) the generation of intensity profiles (Figure 6A). We included a collision detection process during the traverse algorithm to eliminate duplicated data from two connected lipid droplets (see Methods). We manually picked lipid droplets with distinguishable contact site domains and defined the size of the domains by measuring the distance between local intensity minima (Figure 6A, bottom right). The average size of lipid droplets we analyzed for ER-LD and mito-LD contact sites is similar (Figure 6B), ruling out lipid droplets’ size as a contributing factor to the following analyses. The minimal size of contact sites from this analysis is 0.237 µm (Figure 6C, in mito-LD group), which is consistent with the lateral resolution of confocal microscopy.

**Figure 6.**
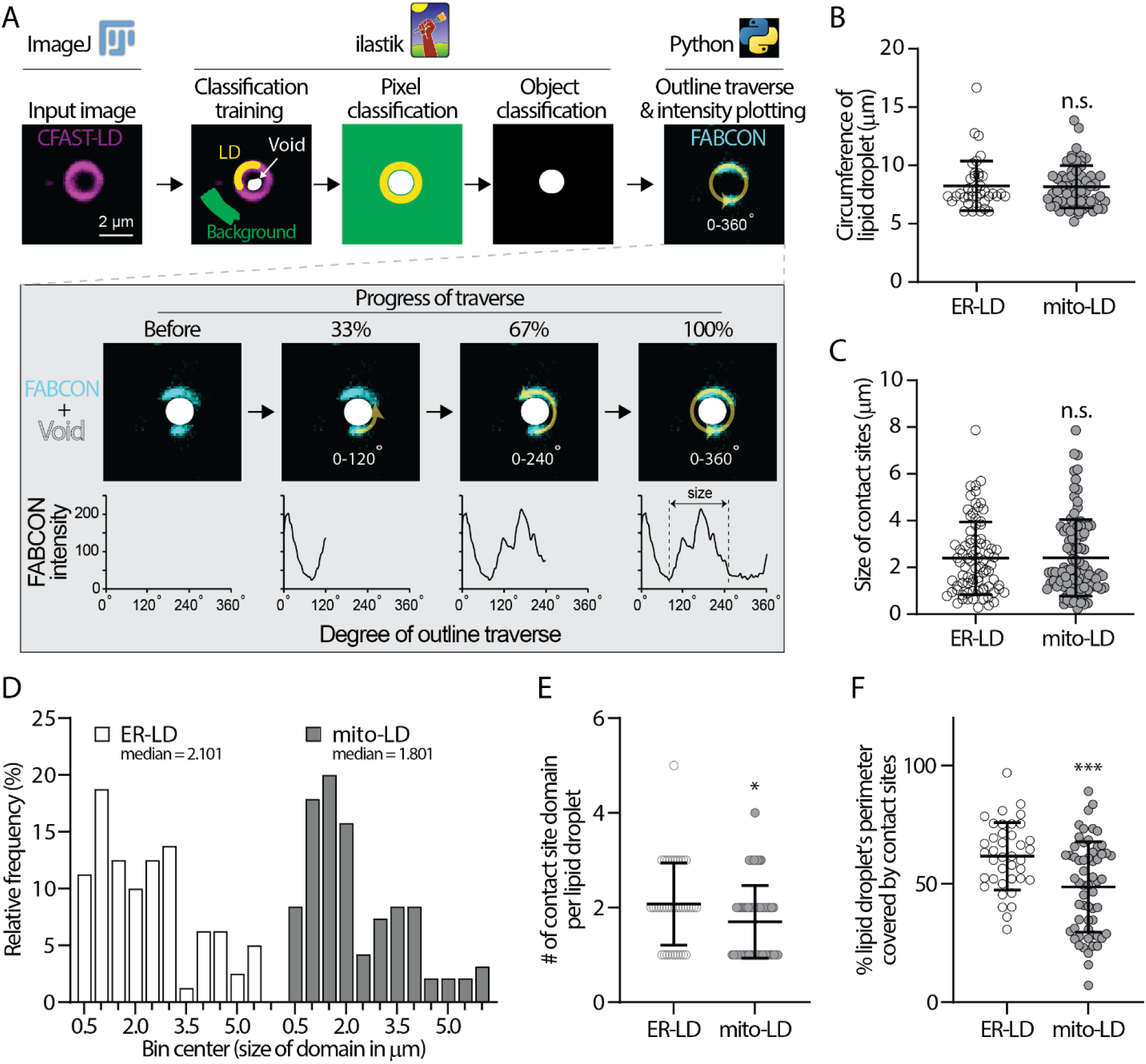
COSIMA analysis of ER-LD and mito-LD contact sites. (A) A flowchart describing the COSIMA pipeline. (B) Circumference of lipid droplets analyzed by COSIMA in ER-LD and mito-LD group. Raw data and mean ± standard deviation are shown (39-60 lipid droplets). (C) Size of ER-LD and mito-LD contact sites measured by COSIMA. Raw data and mean ± standard deviation are shown (81-99 contact sites). (D) Histogram of domain size of ER-LD and mito-LD contact sites. (E) Number of ER-LD and mito-LD contact sites on each lipid droplet. Raw data and mean ± standard deviation are shown (39-60 lipid droplets). (F) Fraction of lipid droplet’s perimeter covered by ER-LD and mito-LD contact sites. Raw data and mean ± standard deviation are shown (39-60 lipid droplets).

After analyzing more than 80 domains from each group, we found that the domain size of ER-LD ranges from 0.291 to 7.873 µm with a mean of 2.392 µm (Figure 6C). Similar range of domain size was found in mito-LD group: 0.237 to 7.861 µm with a mean of 2.410 µm. Interestingly, the median size of mito-LD (1.801 µm) and ER-LD contact sites (2.101 µm) were smaller than their corresponding means. When using the mean as a central tendency of the population, 63% of the mito-LD and 57% ER-LD have domain sizes below their respective means (data not shown). All normality tests rejected normal population distribution of contact site length from the two groups (Table 3). Histograms of domain size from both contact sites were indicative of multimodal distribution. Specifically, most ER-LD population was distributed between 0.5 to 3.5 µm with a smaller subpopulation between 4 to 5.5 µm (Figure 6D). The divide of mito-LD population appeared to be at 2.5 µm with one subpopulation ranging from 0.5 to 2.5 µm and the other from 3 to 6 µm. These results demonstrated that the size of individual contact sites are heterogenous and suggested distinct spatial determinants for each contact site subpopulation.

**Table 3.** Test for normal distribution.

| Test | mito-LD | ER-LD |
| --- | --- | --- |
| D'Agostino & Pearson test |  |  |
| P value | <0.0001 | 0.0015 |
| Passed normality test (alpha=0.05)? | No | No |
| Anderson-Darling test |  |  |
| P value | <0.0001 | 0.0002 |
| Passed normality test (alpha=0.05)? | No | No |
| Shapiro-Wilk test |  |  |
| P value | <0.0001 | 0.0001 |
| Passed normality test (alpha=0.05)? | No | No |
| Kolmogorov-Smirnov test |  |  |
| P value | <0.0001 | 0.0204 |
| Passed normality test (alpha=0.05)? | No | No |

In addition, each lipid droplet harbored an average 2.07 ER-LD contact sites which covered 61.7% of lipid droplet’s circumference (Figures 6E-F). In contrast, mito-LD contact sites only occupied 48.75% of the perimeter of a lipid droplet (Figure 6F), which was primarily due to only 1.70 mito-LD contact sites per lipid droplet (Figure 6E). Interestingly, contact sites of various sizes can exist on the same lipid droplets (Figures 4B, 5B, 6A, and 7A). Overall, these analyses are consistent with the observation that ER-LD contact sites are more prevalent on lipid droplets than mito-LD contact sites (Table 2).

### FABCON is generalizable to visualize organelle contact sites

To demonstrate the generalizability of FABCON, we focused on PX-LD and ER-mito contact sites. As detected by FAB^PX-LD^, PX-LD was the least abundant contact sites of lipid droplets. In U2OS cells, only 66% of lipid droplets could make contact sites with peroxisomes; among these lipid droplets, 93% displayed partial coverage on lipid droplets (Figure 7A, Table 2). When FAB^PX-LD^ was produced in Spastin KO cells, a moderately reduction was detected for PX-LD contact sites (Figure 7B), consisted with the role of Spastin as a tether protein for PX-LD contact sites (Chang et al., 2019). Our data also suggested additional PX-LD tethering mechanisms exist. In contrast to the low occurrence of PX-LD contact sites, ER-mito contact sites revealed by FAB^ER-mito^ (implemented with CFAST10 of lower affinity) (Tebo and Gautier, 2019) were abundant and distributed throughout the entire cell (Figure 7C). ER-mito contact sites appeared to be distinct foci of various size and intensity along mitochondria (Figures 7C inset; 7D). In summary, our data demonstrated that FABCON is readily applicable to detect any contact sites of interest providing proper organelle targeting motif.

**Figure 7.**
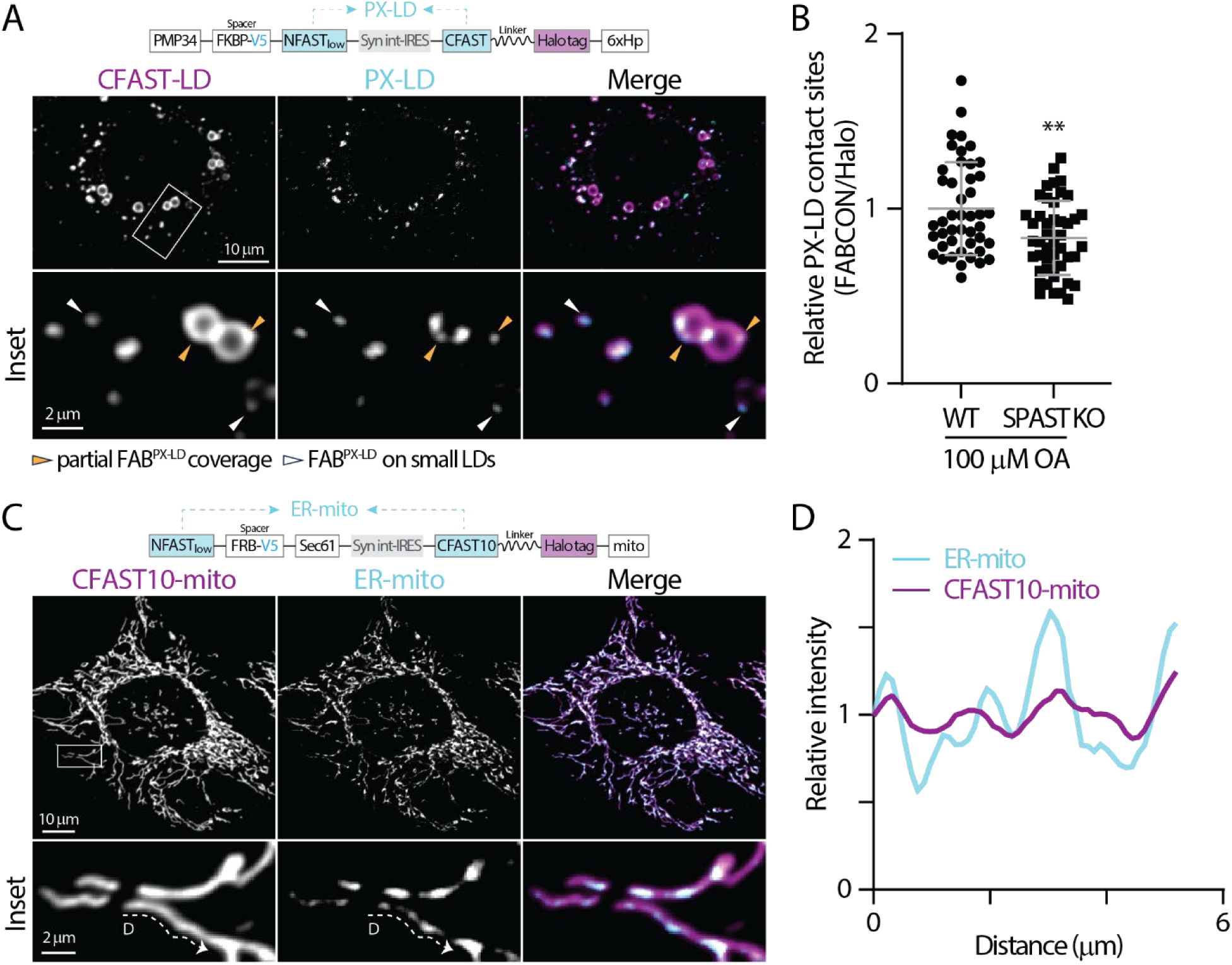
Detection of PX-LD and ER-mito contact sites using FABCON. (A) Detection of lipid droplets and PX-LD contact sites in OA-treated U2OS cells producing FAB^PX-LD^ (top) monitored by confocal microscopy. Representative maximal intensity projected (MIP) images from three axial slices (∼1 µm in total thickness) are shown. (B) Relative levels of PX-LD contact sites in WT and SPAST KO U2OS cells. Raw data and mean ± standard deviation are shown (45 cells for each condition from three independent experiments). (C) Detection of mitochondria and ER-mito contact sites in OA-treated HeLa cells producing FAB^ER-mito^ (top) monitored by confocal microscopy. Representative MIP images from 3 axial slices (∼1 µm in total thickness) are shown. (D) Intensity profiles of CFAST10-mito and ER-LD from the dashed line in C.

## Discussion

Organelle contact sites are an important subcellular architecture of functional integration by providing a platform for inter-organelle transport of a variety of biomolecules, positioning organelles, and facilitating organelle morphogenesis (Abrisch et al., 2020; Lee et al., 2020; Prinz et al., 2020). Since the initial observations of contact sites via EM in 1950s (Palade, 1952; Porter and Palade, 1957), significant progress has been made in uncovering the mechanistic and functional insights into these minute foci. Nonetheless, determining how these nanoscale subcellular foci are dynamically regulated remains challenging due to the limited spatial-temporal resolution of imaging technologies. During the past decade, proximity-induced reporters, such as split FP (Cieri et al., 2018; Eisenberg-Bord et al., 2016; Harmon et al., 2017; Kakimoto et al., 2018; Shai et al., 2018), dimerized-dependent FP (ddFP) (Alford et al., 2012; Miner et al., 2023a), and fluorescence resonance energy transfer FP pairs (Csordas et al., 2010; Naon et al., 2016; Poteser et al., 2016; Venditti et al., 2019; Wong et al., 2018), have been applied to probe organelle proximity at contact sites. Though these approaches are generalizable and straightforward to apply, the implementation has encountered many roadblocks, including irreversibility, fluorescence leakiness, and low signal-to-noise readout. We aimed to use splitFAST to implement this proximity-induced reporter approach to circumvent these difficulties. After vigorous engineering and validation using synthetic biology, LM and EM, and basic biochemistry, we successfully implemented FABCON to dynamically detect ER-LD, mito-LD, PX-LD, and ER-mito contact sites. FABCON’s overall advantages include reversibility, low fluorescence background, and straightforward quantification based on FABCON intensity.

We had to first overcome the challenging issue of lipid droplet targeting by engineering the synthetic 6xHp based on Spastin’s lipid droplet inserting hp motif (Chang et al., 2019). 6xHp displayed significant enrichment on lipid droplets and minimal ER distribution upon expression at moderate levels, which greatly facilitated the implementation of FABCON. We envision that 6xHp will allow us to bring other reporters onto the surface of lipid droplets for additional applications such as proximity labeling-proteomics. Though 6xHp minimally affected lipid droplets’ functions when transiently introduced into cells, the predicted molecular weight of 6xHp is over 35 kDa and it is expected to be highly hydrophobic (data not shown). Further optimization is needed to solve these limitations.

Another unexpected engineering issue is related to the affinity of splitFAST. To our surprise, splitFAST_high_ of sub-µM *K*_d_ significantly altered organelle distribution and interaction while splitFAST_low_ of 220 µM *K*_d_ completely reversed the effects on organelles to resting state. Confining splitFAST to 2-dimensional membranes could substantially enhance their effective *K*_d_ at organelle contact sites. Nonetheless, compared to rapamycin-induced FRB-FKBP dimerization with a *K*_d_ at sub-nM range (Banaszynski et al., 2005), the affinity of splitFAST_high_ self-complementation was rather weak, yet strong enough to drive contact sites formation, suggesting that bimolecular complementation of a sub-µM *K*_d_ reporter at organelle contact sites is practically irreversible. We speculated that the irreversible traditional split FPs at contact sites would significantly remodel organelle distribution, leading to cellular stress and/or adaptation. Another affinity related issue is the stabilization of NFAST-CFAST complementation upon the addition of fluorogen, which can potentially lead to contact sites formation (Rakotoarison et al., 2023; Tebo and Gautier, 2019). To circumvent the stabilization issue, we used a low concentration (<3 µM) of fluorogen and only performed imaging experiments shortly after the addition. Experiments with higher fluorogen concentrations or with incubation periods over 30 min are not recommended. Overall, our observations suggest that organelle contact sites are likely maintained and subjected to rapid turnover via a collection of weak endogenous tethers. The affinity of organelle interaction at contact sites remains an open question. Further work with the pairs of splitFAST with a wide range of self-complementation affinity (Rakotoarison et al., 2023) will help provide needed quantitative information about the affinity of organelle tethering at contact sites.

The reversibility of FABCON provides a wider temporal window to monitor the dynamics of organelle contact sites upon stimulation. We demonstrated that both ER-LD and mito-LD contact sites are dynamically upregulated upon distinct manipulation at cellular level. We envision that FABCON can be used as a reliable readout to address the mechanisms behind these regulations. In addition, FABCON revealed the heterogeneity of contact sites at single lipid droplet level, which will help answer questions related to an organelle’s ability to form contact sites formation and to the subcellular distribution of individual contact sites. Furthermore, our data demonstrated that FABCON can provide intensity-based readout for the levels of contact sites, indicating the possibility of large-scale screening using flow cytometry.

Built on the current understanding that ER-LD contact sites are important for lipid droplet biogenesis (Hugenroth and Bohnert, 2020), FAB^ER-LD^ further revealed the temporal correlation of these sites during this process. We speculate that ER-LD contact sites provide a spatial platform for lipid and/or fatty acid trafficking to fuel the initial growth of lipid droplets, and the ER association may need to be relaxed for further lipid droplet expansion. The negative correlation between seipin and ER-LD contact sites is another interesting observation. Seipin KO cells showed minimal defects in total lipid droplet content (Salo et al., 2019; Wang et al., 2016). Perhaps ER-LD contact sites provided a compensatory mechanism for lipid droplet biogenesis in seipin-deficient conditions.

The association between mitochondria and lipid droplets is expected to play an important role in mitochondrial FAO and cellular energetics. While many reports have shown a correlation between mito-LD contact sites and fatty acid trafficking and oxidation (Rambold et al., 2015; Sadh et al., 2017; Talari et al., 2023; Wang et al., 2021), others have demonstrated that lipid droplet-associated mitochondria have low FAO activity (Najt et al., 2023). Our data from FAB^mito-LD^ may help explain this discrepancy from a temporal point of view. Although mito-LD contact sites are generally upregulated following isoproterenol and 2DG treatment, the extents of upregulation during these conditions are markedly distinct. This temporal information will help guide future mechanistic studies. Further investigation of how mito-LD contact sites are regulated in physiological-relevant cell types, such as hepatocytes, muscle cells, and adipocytes, could shed light on the contribution of these sites in cellular energetics.

To systematically analyze FABCON-derived data of higher spatial precision and acquire statistically meaningful information, we created an automated line scanning pipeline, COSIMA, based on open-source software and codes. COSIMA revealed that the size of contact site domains are heterogenous; contact sites of different lengths could be found on the same lipid droplets. This observation raised possibilities such as: (i) contact sites of different size are maintained via a distinct set of tethers and likely create functional segregation and/or (ii) the size reflects the available lipid droplet surface for contact sites due to physical confinements. Our approach combined with other manipulations may provide insights into lipid droplet-organelle contact sites heterogeneity. With reasonable modifications and optimizations, this pipeline can be readily applied for line scanning on the perimeter of any enclosed objects, such as individual mitochondria or phase condensates.

We envision the next phase for the FABCON development is the generation of orthogonal splitFAST reporters to detect multiple contact sites simultaneously. We did not explore green- and red-splitFAST because they appear to have comparable affinity as splitFAST_high_ (Tebo et al., 2021). We have tried split UnaG, another reversible fluorogenic split reporter (To et al., 2016), but did not observe signal at contact sites (data not shown). One possibility is to utilize split Halo tag and recently developed reversible Halo ligands (Ishikawa et al., 2012; Kompa et al., 2023; Minner-Meinen et al., 2021; Shao et al., 2021) to develop orthogonal split FPs that work with splitFAST. Additionally, it is possible to combine FABCON with other ddFP-based tool kits to investigate multiple contact sites (Alford et al., 2012; Miner et al., 2023a). Overall, our work provides a new tool kit for the detection of lipid droplet-organelle contact sites and beyond; we have demonstrated that FABCON is capable of revealing the dynamics of contact sites and generating observation-based hypotheses.

## Methods

### Cell lines and transient transfection

HeLa, U2OS and HepG2 cells were purchased from American Type Culture Collection (ATCC). All cell lines were maintained in media following vendors’ recommendations. VAP DKO and parental HeLa cells were provided by Dr. Wade Harper at Harvard Medical School and were grown in Eagle’s minimal essential medium (EMEM, Thermo Fisher Scientific, Waltham, MA, USA) with 10% fetal bovine serum (FBS) and 1X penicillin/streptomycin solution (Corning, Corning, NY, USA). Seipin KO and wildtype SUM159 cells were obtained from the Farese and Walther lab at the Memorial Sloan Kettering Cancer Center. SUM159 cells were cultured in DMEM/F-12 GlutaMAX (Life Technologies, Carlsbad, CA, USA), supplemented with 5% FBS, 1X penicillin/streptomycin solution, 1 µg/mL hydrocortisone (Sigma-Aldrich, St. Louis, MO, USA), 5 µg/mL insulin (Sigma-Aldrich), and 10 mM HEPES (pH 7.0) (Sigma-Aldrich). Transient transfection was performed using TransIT-LT1 (Muris Bio LLC, Madison, WI, USA) or Lipofectamine 3000 (Thermo Fisher Scientific) according to manufacturers’ instructions for 16– 20 h. To knock down PLIN 5, HeLa cells were transfected with 25 nM scramble or PLIN 5 siRNA (Dharmacon, Lafayette, CO, USA) using TransIT-TKO (Mirus Bio LLC) according to manufacturer’s instructions.

### DNA Plasmids

FABCON plasmids were generated as the following. We first generated plasmids containing FP- (mApple, mCherry, or Halo) tagged organelle targeting motifs. FP-1xHp was generated by inserting a 1xHp PCR fragment into an FP-C1 vector using the EcoRI and BamHI restriction sites. FP-6xHp was created by replacing 1xHp in FP-1xHp with a 6xHp gBlock using the BsrGI and BamHI restriction sites. PMP34-FP was generated by inserting a PMP34 PCR fragment into an FP-C1 vector using the NheI and AgeI restriction sites. A CFAST gBlock or an annealing nucleotide and NFAST PCR fragment were inserted into FP-6xHp using the NheI and AgeI restriction sites to generate CFAST-FP-6xHp and NFAST-FP-6xHp, respectively. A linker gBlock was then inserted into CFAST-FP-6xHp using the AgeI restriction site to generate CFAST-linker-FP-6xHp. CFAST-FP-ER was created by replacing 6xHp in CFAST-FP-6xHp with a gBlock containing an ER targeting motif (Cho et al., 2020) using the SacI and BamHI restriction sites. NFAST-FP-ER, NFAST-FP-Sec61β, and NFAST-FP-mito were created by replacing 6xHp in NFAST-FP-6xHp with gBlocks containing an ER targeting motif, Sec61β, and mitochondria targeting motif (Benedetti et al., 2020), respectively, using SacI and BamHI. To generate PMP34-FP-NFAST, we inserted an NFAST fragment into PMP34-FP using the Kpn2I and NotI restriction sites. The Halo-TNFa-RUSH construct (Weigel et al., 2021) was used as a backbone to generate bicistronic IRES plasmids. In brief, (5’–3’) NFAST-containing PCR fragments (NheI-NotI digested), a synthetic intron-IRES (NotI-SgsI digested) fragment, and the PCR product of CFAST-linker-Halo-6xHp (SgsI-Acc65I digested) were ligated into the backbone digested with NheI and BsrGI restriction enzymes. To replace FP with spacers in all NFAST-containing halves in IRES plasmids, gBlocks of various spacers (see Figure S4) were inserted into PCR-amplified backbones omitting FP by InFusion cloning. Lentiviral plasmids were generated by inserting PCR fragments containing coding sequences of NFAST- and CFAST-containing halves flanking IRES element into the lentiviral SJL12 backbone (provided by the St. Jude Vector Core, Memphis, TN, USA) digested with AgeI and NotI restriction enzymes using InFusion cloning.

APEX2-6xHp was generated by replacing mApple in mApple-6xHp with APEX2 PCR fragments using the NheI and EcoRI restriction sites. mEmerald-Sec61β (#90992) and Halo-TNFα-RUSH (#166901) were obtained from Addgene (Watertown, MA, USA). All plasmids were confirmed via sequencing. All oligonucleotides (Table S2) and gBlocks were synthesized by Integrated DNA Technologies (Coralville, IA, USA).

### Lentivirus production

Production and titration of lentiviral vectors were performed as described previously (Bauler et al., 2020). Briefly, SJ293TS cells were transfected with the transfer vector and the helper plasmids pCAG-kGP1-1R, pCAG-VSVG and pCAG4-RTR2 using PEIpro (Polyplus Transfection, Strasbourg, France) and grown in Freestyle 293 Expression media (Thermo Fisher Scientific) at 37°C with 8% CO_2_ and shaking at 125 RPM. The next day, Benzonase (Millipore-Sigma, Burlington, MA, USA) was added to the transfected cells with a final concentration of 6.25 U per mL. Vector supernatants were collected 48 hours post-transfection, clarified by centrifugation at 330 × g for 5 minutes and 0.22 µm filtered. Lentiviral vector containing supernatants were adjusted to 300 mM NaCl, 50 mM Tris pH 8.0 and loaded onto an Acrodisc Mustang Q membrane (Pall Life Sciences, NY, USA) according to the manufacturer’s instructions using an Akta Avant chromatography system (GE Healthcare Bio-Sciences, Pittsburgh, PA, USA). After washing the column with 10 volumes of 300 mM NaCl, 50 mM Tris pH 8.0, viral particles were eluted from the column using 2 M NaCl, Tris pH 8.0. Viral particles were formulated into either X-VIVO™ 10 or X-VIVO™ 15media (Lonza, Walkersville, MD, USA) or phosphate buffered saline containing 1% human serum albumin (Grifols Biologicals, Los Angeles, CA, USA) using either a PD10 desalting column (GE Healthcare) or a Vivaflow 50 cassette (Sartorius) according to the manufacturer’s instructions to achieve an approximate 50-fold concentration from the starting material, 0.22 µm sterile filtered, aliquoted and stored at −80°C.

Titration of lentiviral vectors was performed by transducing HOS cells (ATCC CRL-1543) with serially diluted vector preparations in the presence Polybrene (5-8 µg/mL; Millipore Sigma, Burlington, MA, USA). HOS cells were grown in DMEM (Corning, Corning, NY, USA) supplemented with 10% fetal bovine serum (Seradigm, Radnor, PA, USA) and 2 mM L-alanyl-L-glutamine (Corning) at 37°C with 5% CO_2_. Four days post-transduction, genomic DNA was isolated from transduced HOS cells using a Quick-DNA Miniprep kit (Zymo Research, Irvine, CA, USA). Vector titers were determined by calculating the ratio between the copies of HIV psi and every two copies of RPP30 via QX200 digital droplet PCR system (Bio-Rad, Carlsbad, CA, USA), multiplied by the number of cells transduced and if necessary, multiplied by the dilution factor.

### Fluorescence Microscopy Imaging

#### General imaging and FRAP

All cells were grown and transiently transfected or infected with lentiviral vectors on Lab-Tek II chambered #1.5 coverglasses (Thermo Fisher Scientific) or MatTek dishes with #1.5 coverslip (MetTek, Ashland, MA, USA). Prior to imaging, media were replaced with FluoroBrite DMEM (Thermo Fisher Scientific) supplemented with 5% FBS and penicillin/streptomycin and imaged at 37°C. Confocal microscopy was conducted on a custom-built Nikon microscope equipped with a Yokogawa CSU-W1 Spinning Disk unit, XY galvo scanning module, and Tokai Hit STXG CO_2_ incubation system using 60x (CFI APO TIRF, NA 1.49 oil) and 10x (CFI Super Fluor, NA 0.50) objectives. Photobleaching experiments were performed on a Nikon microscope using XY galvo scanning module. Structured illumination microscopy (Scorrano et al.) imaging was performed using the Plan Apochromat 63×/1.4 oil objective on a Zeiss ELYRA7 Microscope.

#### FABCON imaging

Cells were seeded on an 8-well coverglasses at a density of 2,000 to 4,000 cells per well for 24 h followed by FABCON lentivirus infection at a multiplicity of infection (MOI) of 300 for FAB^ER-LD^ and FAB^mito-LD^; MOI of 500 for FAB^PX-LD^ unless otherwise indicated. After 24 h of infection, the infection media was replaced with fresh culture media with indicated amount of oleic acid (OA). The next day, infected cells were imaged in the presence of 100 nM of Halo ligand JF646, which labels Halo tagged 6xHp on lipid droplets. Then, 3 µM of HBR-2,5DOM (Mineev et al., 2021) was added upon imaging acquisition to label contact sites FABCON, which was imaged with a 488-nm excitation and a 560/50 emission filter. For intensity-based analysis, all images were taken within 10–15 min following fluorogen addition. It is important to note that a maximum of 3 µM fluorogen should be applied to circumvent issues related to enhancing contact site formation.

For ER-LD contact site imaging during LD biogenesis, HeLa cells producing FAB^ER-LD^ were incubated with 500 µM OA and imaged at indicated time points. To evaluate the effect of Seipin on ER-LD contact sites, seipin KO and wild-type SUM159 cells were infected with lentivirus producing FAB^ER-LD^ at an MOI of 500. Following 24 h of viral transduction, the medium was replaced with fresh medium supplemented with 20 µM or 100 µM OA. The cells were then imaged for the ER-LD contact sites the next day. To examine how isoproterenol and 2DG affect mito-LD contact sites, HeLa cells were seeded and transduced with lentivirus producing FAB^mito-LD^. Following 24 h of viral transduction, the medium was replaced with fresh EMEM media supplemented with 100 µM oleic acid. The next day, prior to imaging, media were replaced with imaging media without (control) and with 4 mM 2DG or 10 µM isoproterenol. Cells were imaged at indicated time points.

#### Lattice light sheet microscopy (LLSM)

U2OS cells were seeded on CS-5R coverslips (Multi Channel Systems, Reutlingen, BW, Germany) and infected with lentivirus producing FAB^ER-LD^. Twenty-four hours later, the cells were replenished with fresh DMEM/F12 medium supplemented with 100 μM of oleic acid and incubated overnight. U2OS cells were then imaged in FluoroBrite DMEM with 5% FBS and 100 nM JF646 Halo ligand. Live imaging was acquired with a lattice light sheet microscope (Intelligent Imaging Innovations, Denver, CO, USA) operating in single camera mode using a custom emission filter (Chroma ZET488/640m-TRF). Excitation of HBR-2,5DOM and JF646 Halo ligand was achieved using 488nm and 640nm lasers (MPB Communications Inc.) respectively. The desired excitation profile was achieved using a multi-bessel beam interference pattern ((Liu et al., 2023), crop factor ε = -.15) filtered through an annular mask with inner and outer NA; 0.472 and 0.55; to create a light sheet with empirical propagation length and axial thickness approximately 20µm and 1.05µm at 488nm, respectively. The propagation length value is obtained from an image of a single-Bessel illumination pattern emitted by fluorescein dye solution excited at 488nm and is the full width at half maximum (FWHM) of the intensity profile along the direction of propagation at the maximum intensity. Note that the length of this single-Bessel beam, created by fully illuminating the annular mask, is a slight under-estimate of the multi-bessel beam produced by the patterned illumination at the same mask. The axial thickness value is obtained from the measured XZ PSF, captured by imaging the emission of a fluorescent 100 nm microbead scanned through the illumination pattern, and is the FWHM of the profile along the axial direction at a pattern maximum. This is well matched to the theoretical values of propagation length (22.5 µm) and axial thickness (1.0 µm) for this pattern at 488 nm (Shi et al., 2022). Emission was detected with a Zeiss 1.1NA, 20x water immersion detection objective. Volume images were acquired in sample scanning mode with a 0.3-µm step l. Raw images were subsequently deconvolved using a standard Lucy-Richardson algorithm with experimentally acquired lattice light sheet point square functions and resampled to remove the skew orientation introduced by sample scanning.

### Cell fixation and immunostaining

All procedures were performed at room temperature and all washing steps were done using Dulbecco’s Phosphate-Buffered Saline (DPBS) for 5 min unless otherwise indicated. Cells were rinsed with DPBS and fixed with 4% paraformaldehyde and 0.1% glutaraldehyde in DPBS for 20 min. Fixed cells were quenched with 100 mM glycine in DPBS, washed twice, and permeabilized by 0.3% Triton X-100 in DPBS for 20 min. Permeabilized cells were then blocked with 5% normal donkey or goat serum in DPBS for 1 h followed by incubation with primary antibody (anti-V5 antibody, CST, cat# 13202S; anti-PLIN 2, Abcepta, cat# AP5118c) in DPBS with 1% BSA at 4°C overnight. After three washes, the samples were incubated with fluorescent secondary antibody (1: 2,000 in dilution) for 1 h. The stained samples were washed three times and imaged with confocal microscopy at room temperature.

### Image Analysis

All image analyses were performed using Fiji (Schindelin et al., 2012) (National Institutes of Health) unless otherwise indicated. All intensity analyses were subjected to background subtraction. To obtain relative intensity profiles, background-subtracted intensity values from different conditions were normalized to that at the first time point or in control groups. To measure relative contact sites level, regions of interest (ROIs) of lipid droplets were generated by thresholding CFAST-LD images. These ROIs were then applied to FABCON and CFAST-LD images to obtain their mean gray values. The relative level of contact sites was obtained by normalizing the FABCON mean gray value to that of Halo-6xHp.

For colocalization analysis, the machine learning software ilastik (version 1.4) (Berg et al., 2019) was trained to segment mitochondria (mito), lipid droplets (LD), and peroxisomes (PX) from Maximum Intensity Projected (MIP) images to create binary masks. The masks were then used as input for JACoP (Just Another Colocalization Plugin) in Fiji to acquire Pearson’s coefficient between two organelles.

For measuring relative lipid droplet content, HeLa cells transiently transfected with Halo-6xHp (stained with JF646) were fixed after indicated treatment. Fixed cells were post-stained with BODIPY and confocal images of 6xHp and lipid droplets were acquired using a 10x objective. Background staining of BODIPY throughout entire cytoplasm was threshold to generate ROIs for single cells using Wand (tracing) tool in Fiji. These ROIs were then applied to 6xHp and lipid droplets images to obtain their mean gray values. Corresponding data of 6xHp and lipid droplets were binned based on 6xHp’s values: (i) <20 is considered absence of expression (-), (ii) 20-100 represents low expression (+), and (iii) 200-500 indicates moderate 6xHp expression (++).

### COSIMA

The COntact SIte MApping (COSIMA) was developed to automate the quantification and plotting of fluorescence intensities at contact sites (i.e., ER-LD, mito-LD) based on the segmentation masks created by ilastik. Original *.nd2 files Z-stacks images were converted into maximum intensity projected *.tif images (MIP) using Fiji. For consistency, the MIP was centered around the structure in focus and included an identical number of slices above and below the focus plane in each channel. Ten MIP images of lipid droplets labeled by CFAST-LD were randomly chosen and manually annotated to be used as a training set for a pixel classifier using ilastik. Three predicted classes were evaluated for each pixel: LD membrane, LD void (center of LD), and background. The resulting prediction maps (matched to their original images) were further manually annotated and used to create an object classifier using ilastik. This step improved the accuracy of the detection of LD voids by removing false positive objects incorrectly identified by the pixel classifier. The final output was a binary mask of the LD voids and corresponding features saved in *.tif and *.csv format, respectively. Following critical manual review of the segmentation results on the training dataset, this pipeline was applied to the rest of the images.

COSIMA was created based on the Python programming language and ported to Napari (v 0.4.18, Ahlers, 2023) as a plugin. The software identified each segmented LD void from the input image and created an outline of a defined thickness starting at the edge of the LD void and encompassing the membrane of the LD. The growth of each outline was done sequentially in 1-pixel wide increments for all the objects present in an image. In case of adversarial growth between neighboring objects, a dedicated collision module prevented the formation of overlapping outlines by limiting their local expansion. For all datasets, the analyzed area was set to be 5 pixels wide (= 514 nm) around the void of each LD and was used to measure fluorescence intensities on all channels. Following automated background subtraction, fluorescent intensities were recorded per pixel for each individual layer and averaged radially over the total number of layers analyzed. Final output included measurement tables, LD-specific plots of fluorescence intensities around each object and annotated images allowing manual review and visual confirmation by the users. This pipeline is available at, https://github.com/stjude/cosima.

### EM imaging

To investigate mitochondria-LD contact sites, HeLa cells were plated in MatTek dishes (cat# P35G-1.5-10-C) at a density of 3000 cells/dish and infected with lentivirus producing FAB^mito-LD^ at MOI of 500 the next day. Uninfected and infected HeLa cells were maintained in EMEM medium for an additional 24 h before being supplemented with 100 μM of oleic acid; cells were allowed to grow overnight. For EM imaging, HeLa cells were pre-fixed in a mixture of 2.5% glutaraldehyde, 2% paraformaldehyde, and 0.1M sucrose in 0.1M Sorenson’s phosphate buffer (PB), pH 7.4 overnight at 4°C. The pre-fixed cells were post-fixed in a mixture of 0.5% OsO_4_ and 0.5% K_4_Fe(CN)_6_ in 0.1M PB, pH 7.4 for 10 min in cold beads. The post-fixed cells were contrasted with 0.5% uranyl acetate overnight at 4°C. Following the contrasting, cells were dehydrated in ascending ethanol series (10%, 30%, 50%, 70%, 80%, 90%, 95%, and 100%), infiltrated in Spurr’s resin (25%, 50%, 75%, and 100%), and embedded in absolute Spurr’s resin overnight. Resin-embedded cells were thermally polymerized for 24 h at 70°C. The coverslip under the MatTek glass bottom dish was detached using glass removal fluid and liquid nitrogen. The polymerized cells were sectioned to a 2 μm depth from the block face using a Leica EM ARTOS 3D ultramicrotome (Leica Microsystems, Wetzlar, Germany) and a diamond knife (DiATOME, Nidau, Switzerland). The last two sequential ultrathin sections (900 μm x 900 μm x 100 nm) were obtained on a formvar/carbon-coated slot grid (2 mm x 1 mm). The ultrathin sections were post-stained with 1% uranyl acetate.

Contrasted ultrathin sections were observed using a STEM detector at 15-KeV and the high-angle annular dark field imaging mode of a Carl Zeiss Gemini 460 field emission scanning electron microscope. Twenty cells per group were randomly selected at 1,000 x magnification for an unbiased quantification of contact sites between endoplasmic reticulum and mitochondria. Images at 5,000 x and 10,000 per cell were acquired in lipid-rich regions. Reagents and supplies from Electron Microscopy Sciences, Hatfield, PA, USA unless indicated otherwise.

### APEX2

To prepare cells for proximity labeling with APEX2 and subsequent electron microscope imaging, 1 x 10^4^ U2OS cells were seeded in MatTek dishes (cat# P35G-1.5-10-C) and transfected with 300 ng of APEX2-6xHp construct. After transfection, U2OS cells were maintained in fresh DMEM/F12 medium supplemented with 100 μM of oleic acid overnight. The U2OS cells were then fixed with 2% glutaraldehyde at room temperature for 5 min, followed by glycine quenching. Subsequently, cells were stained with 1x DAB with 10 mM H_2_O_2_ for 15 min on ice. Steps to process samples for EM were performed as described previously (Martell et al., 2017). Samples sectioned at 70nm on a UC-7 ultramicrotome (Leica) to uncoated copper grids and imaged on a Thermo Scientific TF20 TEM at 80keV and images captured with an AMT (Woburn, MA) NanoSprint15 imaging system.

### Lipid droplet floatation assay

Lipid droplets were isolated from HepG2 cells following a previously described protocol (Bersuker et al., 2018). Briefly, 1.2 x 10^7^ cells were transfected with mApple-1xHp and then incubated with 200 μM oleic acid overnight. Cells were subsequently scraped off the plates in cold PBS buffer and resuspended in 2 mL of HLM buffer (20 mM Tris-HCl, pH 7.4, 1 mM EDTA, pH 8.0). Cell lysis was achieved by passing the cell suspension through 27G needles 40 times, followed by centrifugation at 1000 x g for 10 min at 4°C. Supernatants were mixed with 60% sucrose and overlaid with 5% sucrose and HLM buffer in an SW41i tube (Cat. # 344059). The gradient was centrifuged at 28,000 × g for 30 min and allowed to coast to a stop. After centrifugation, buoyant fractions were separated from the gradients with a tube slicer (Cat. # 303811). Lower fractions were collected from the top of the gradients in 1-mL increments, and membrane pellets were resuspended in 1 mL of HLM buffer. The protein concentration of each fraction was determined using a Nanodrop A280 spectrophotometer, and an equal amount of protein was loaded into an SDS-PAGE gel for subsequent analysis with western blot. mApple tagged proteins were probed with anti-RFP antibody (Abcam, cat # ab124754, 1:2000), perilipin 2 (PLIN 2) was probed with anti-ADFP (Plin2) antibody (abcepta, cat # AP5811c, 1:2000), and ER marker calreticulin was detected with anti-calreticulin antibody (Abcam, cat # ab2907).

### Western blot

To validate knockout cell lines and cells treated with siRNAs, cells were lysed with IP lysis buffer (Thermo Fisher Scientific) supplemented with 1X protease inhibitor cocktail (Thermo Fisher Scientific) on ice for 30 mins. The cell lysates were centrifuged at 18,000 x g for 15 min at 4°C. The supernatant was collected and denatured in Laemmli buffer (with 10% beta-mercaptoethanol) at 95°C for 10 min. Proteins were separated on an SDS-PAGE gel and transferred to a nitrocellulose membrane (BIO-RAD, Hercules, CA, USA). The membrane was blocked with EveryBlot blocking buffer (BIO-RAD) for 1 h, and incubated with a primary antibody (1:1,000) overnight. The membrane was then washed with PBST for 10 min x 3 times and incubated with a corresponding secondary antibody (1:2,500). The membrane was then washed again with PBST for 10 min X 3 times before visualization of protein bands. Chemiluminescence was detected using ChemiDoc Imaging System (BIO-RAD). The following antibodies were used in this study: anti-Seipin antibody (Abnova Cat# H00026580-A02), anti-Plin 5 (Progen Cat# GP31), anti-VAPA (Sigma Cat# HPA009174), anti-VAPB (Sigma Cat# HPA013144), anti-Spastin (Sigma Cat# ABN368), and anti-GAPDH (Cell Signaling Cat# 2118).

## Statistical Analysis

Data were statistically analyzed using a two-tailed *t*-test or one-way analysis of variance with Tukey’s multiple comparisons using GraphPad Prism (GraphPad Software; La Jolla, CA). Data distribution was assumed to be normal unless otherwise indicated. Normal distribution test was performed using Prism. *, p < 0.05; ** p < 0.01; ***, p < 0.001; n.s., not significant. Graphs were generated using GraphPad Prism and Adobe Illustrator (San Jose, CA, USA).

## Acknowledgements

We would like to thank Dr. Luke Lavis at Janelia Research Campus for providing JF Halo ligands, Dr. Wade Harper at Harvard Medical School for providing parental and VAP DKO HeLa cells, Drs. Robert Farese, Jr. and Tobias Walther at Memorial Sloan Kettering Cancer Center for providing parental and Seipin KO SUM159 cells, Vector Core Lab at St. Jude for generating lentiviral particles, Jonathon Klein and Dr. Shondra M. Pruett-Mille for generating SPAST KO U2OS cells, Dr. David J. Solecki for LLSM consultation, Dr. Danny M. D’Amore for scientific editing, Drs. J. Paul Taylor, Joseph Opferman, and Wyatt Beyers for feedback on the manuscript, and Gunda Johnson, Marquetta Nebo, Jennifer Maglisco and Monique Payton for administrative and operational assistance. This work was supported by ALSAC (to C.-L.C.) and the National Institutes of Health (1DP2GM150192 to C.-L.C.). The authors declare no competing financial interests.

## Author Contributions

Xiao Li, Rico Gamuyao, and Chi-Lun Chang performed experiments and analyzed data. Ming-Lu Wu and Alex Carisey designed and created COSIMA pipeline. Woo Jung Cho and Nathan B. Kurtz performed EM sample preparation and imaging and Camenzind G. Robinson supervised the EM experiments. Xiao Li, Sharon V. King, and R.A. Petersen performed LLSM imaging. Xiao Li and Abbas Shirinifard performed 3D rendering of LLSM images. Daniel R. Stabley conceived of LLSM imaging approaches and supervised LLSM imaging and data processing. Caleb Lindow and Leslie Climer generated a subset of plasmids. Francesca Ferrara generated SJL12 lentiviral backbone for cloning and Robert E. Throm supervised the Vector Core Lab for lentivirus production. Alison G. Tebo generated splitFAST_low_ and HBR-2,5DOM and provided critical feedback on the implementation of splitFAST. Chi-Lun Chang, Xiao Li, and Rico Gamuyao wrote the manuscript with the input from all co-authors. Chi-Lun Chang conceived and supervised this project.

## Abbreviations

2DG: 2-deoxy-d-glucose
6xHp: 6x tandem repeats of Hp
BiFC: bimolecular fluorescence complementation
COSIMA: contact site mapping
DMEM: Dulbecco’s Modified Eagle Medium
DPBS: Dulbecco’s Phosphate-Buffered Saline
EM: electron microscopy
ER: endoplasmic reticulum
ER-LD: endoplasmic reticulum-lipid droplet
FABCON: Fluorogen-Activated BiFC at CONtact
FAO: fatty acid oxidation
FAST: fluorescence-activating and absorption shifting tag
FBS: fetal bovine serum
FP: fluorescent protein
FRAP: fluorescence recovery after photobleaching
HBR: hydroxybenzylidene rhodanine
Hp: hairpin
KO: knockout
LLSM: lattice light sheet microscopy
LM: light microscopy
MIP: maximum intensity projected
mito-LD: mitochondria-lipid droplet
MOI: multiplicity of infection
PLIN: perilipin
PX-LD: peroxisomes and lipid droplets
VAP DKO: VAP-A/B double knockout

## Figures and legends

**Figure S1.**
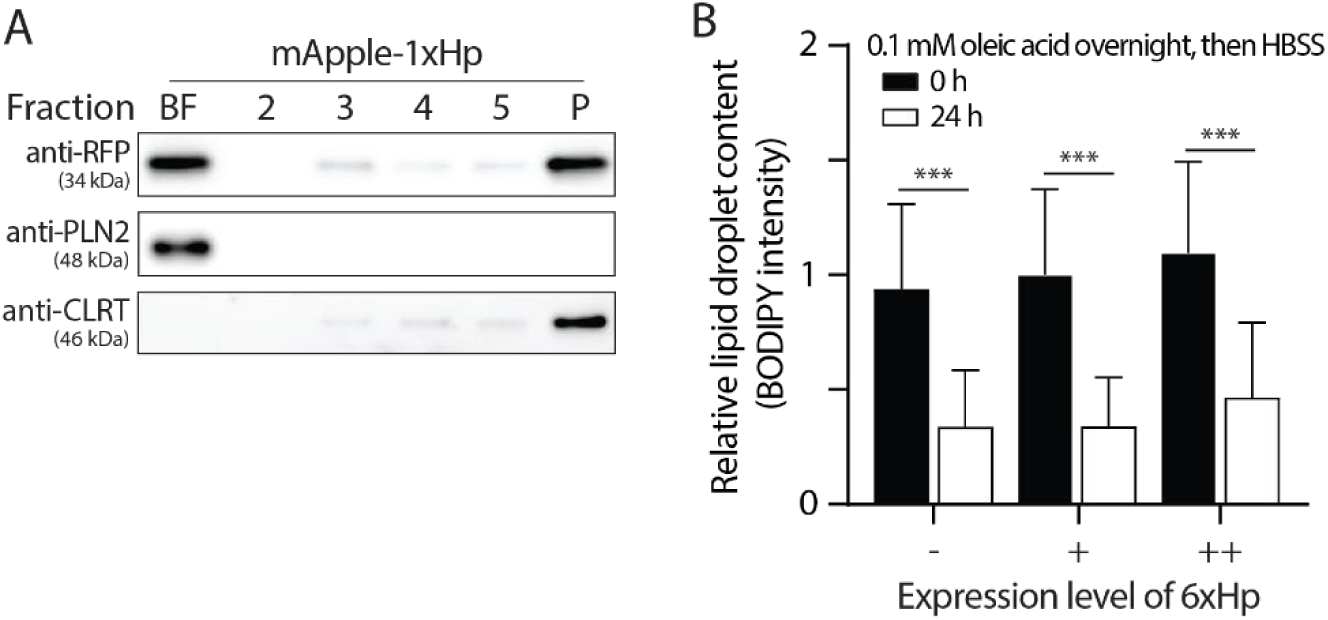
Validation of a synthetic lipid droplet targeting motif. (A) Distribution of overexpressed mApple-1xHp, endogenous perilipin 2 (PLIN2), and endogenous calreticulin (CLRT) in sucrose-gradient cellular fractionations from HepG2 cells treated with 200 µM oleic acid (OA) overnight. BF, buoyant fraction; P, pellet. (B) Relative lipid droplet content in OA-loaded, Halo-6xHp overexpressing HeLa cells incubated with HBSS for 0 or 24 h. Mean ± standard deviation are shown (215-1275 cells from two independent experiments). ‘-‘ indicates absence of 6xHp expression. + and ++ indicate low and moderate expression of 6xHp, respectively.

**Figure S2.**
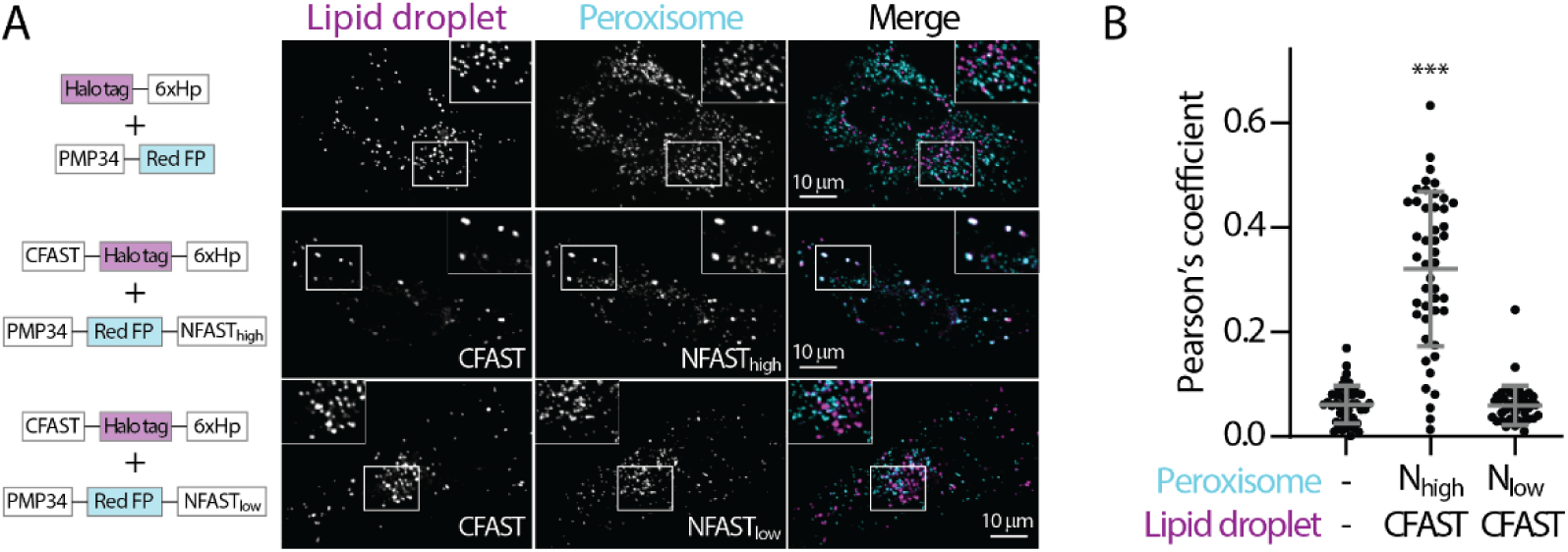
Low affinity splitFAST minimally affects the distribution of lipid droplets and peroxisomes. (A) Distribution of lipid droplet and peroxisome in oleic acid (OA)-treated HeLa cells overexpressing Halo-6xHp and PMP34-mApple (top), CFAST-Halo-6xHp and PMP34-mApple-NFAST_high_ (middle), or CFAST-Halo-6xHp and PMP34-mApple-NFAST_low_ (bottom) monitored by confocal microscopy. Maximal intensity projected images from six axial slices (1.8 µm in total thickness) are shown. (B) Quantification of the Pearson’s colocalization coefficient of lipid droplet and peroxisome described in (A). Raw data and mean ± standard deviation are shown (44-51 cells from three independent experiments).

**Figure S3.**
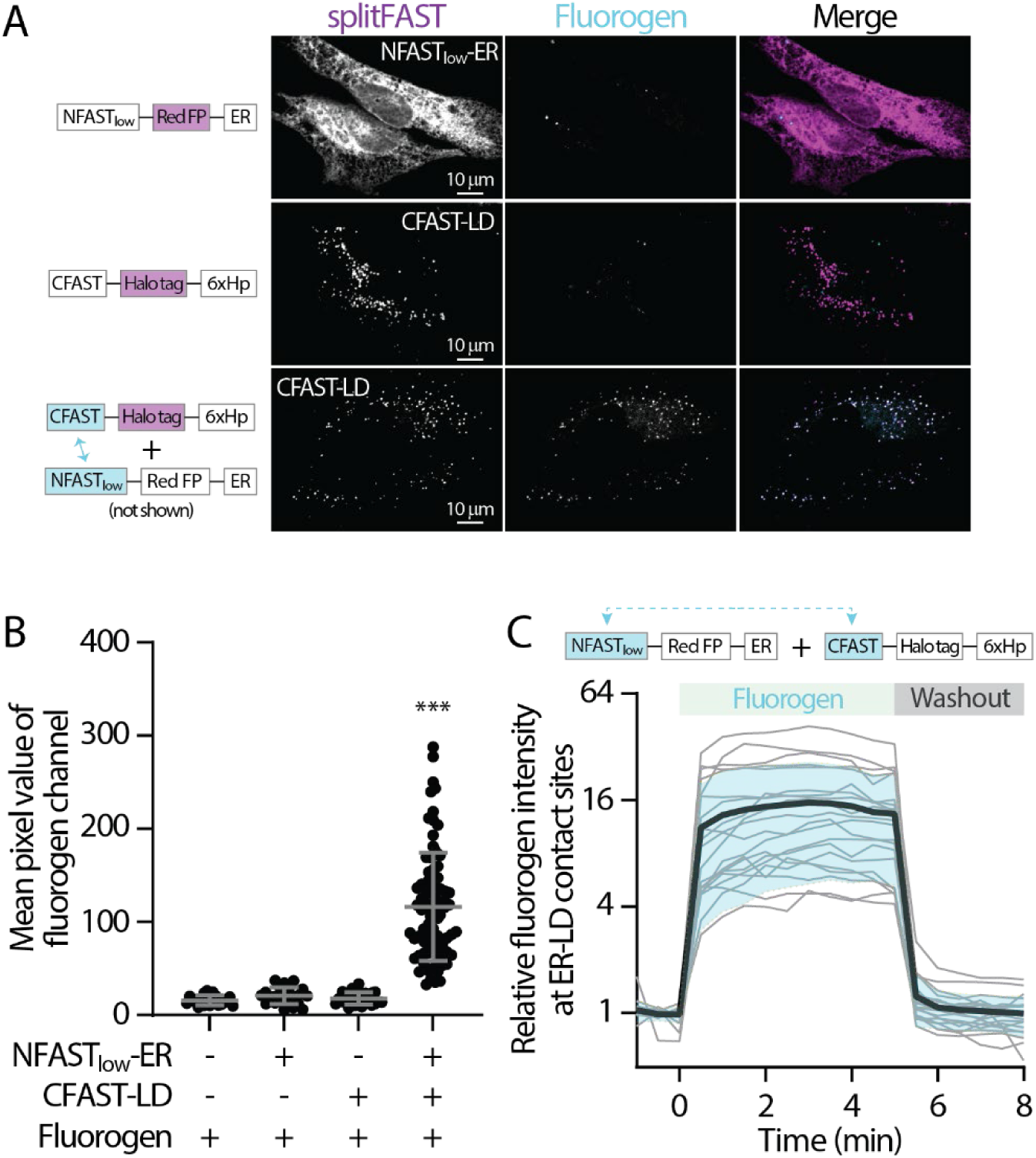
splitFAST remains reversible at organelle contact sites without detectable fluorescence leakiness. (A) Confocal images of HeLa cells expressing NFAST_low_-mApple-ER (top), CFAST-Halo-6xHp (middle), and NFAST_low_-mApple-ER plus CFAST-Halo-6xHp (bottom) in the presence of HBR-2,5DOM. (B) Quantification of fluorogen intensity of (A) and in control HeLa cells. Raw data and mean ± standard deviation are shown (22-82 cells). (C) Relative fluorogen intensity in HeLa cells coexpressing NFAST_low_-mApple-ER and CFAST-Halo-6xHp following HBR-2,5DOM addition and washout monitored by confocal microscopy. Raw data (gray traces) and mean (bold black trace) ± standard deviation (shaded blue) are shown (19-20 cells).

**Figure S4.**
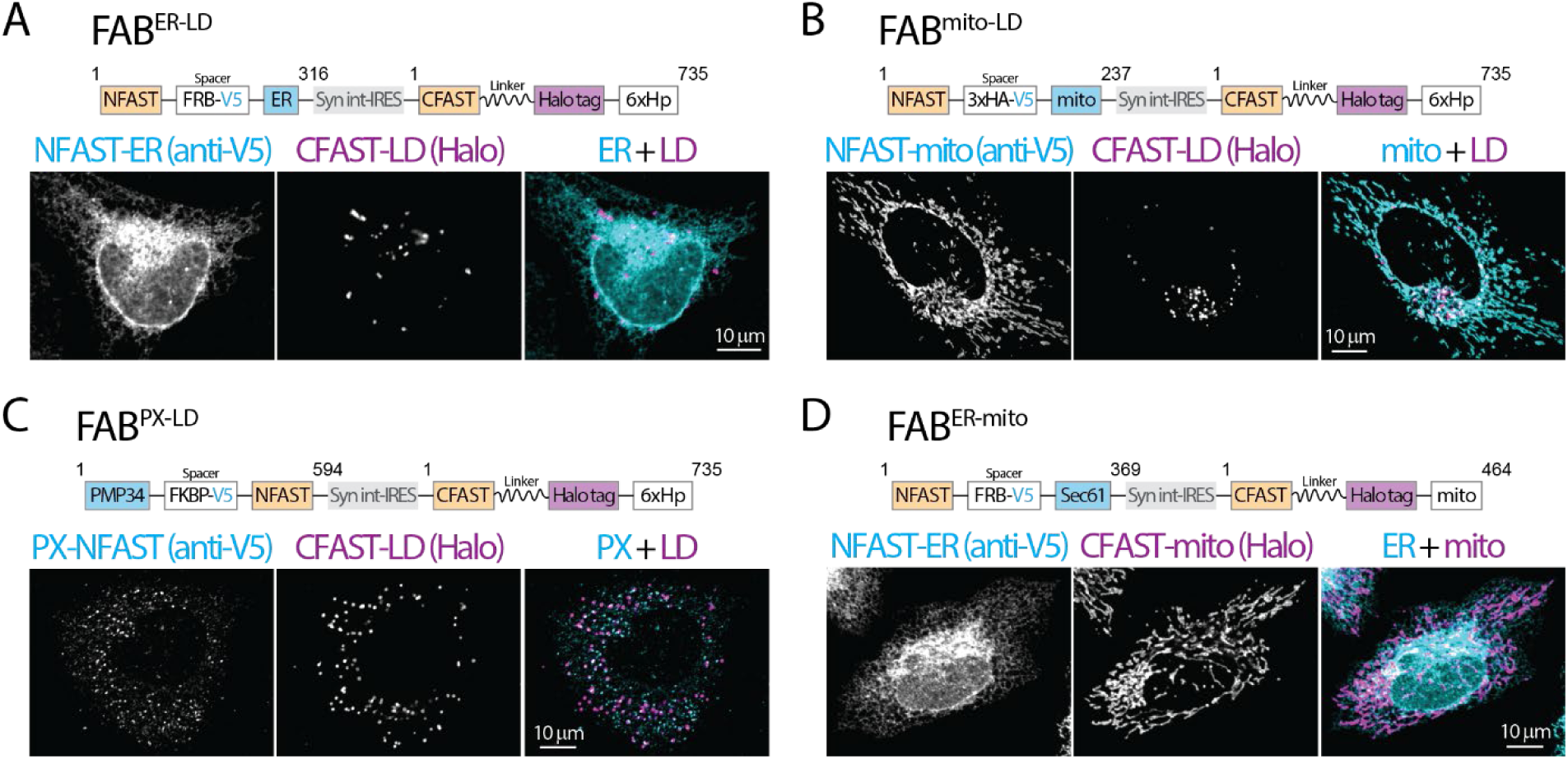
Diagram and validation of FABCON lentiviruses. (A-D) Diagram (top) and organelle distribution of cognate FABCON halves (bottom) of FAB^ER-LD^ (A), FAB^mito-LD^ (B), FAB^PX-LD^ (C), and FAB^ER-mito^ (D) monitor by confocal microscope. NFAST fused organelle marker are immunostained with anti-V5 antibody. Maximal intensity projected images from three axial slices (∼1 µm in total thickness) are shown. Numbers of amino acids are indicated. Syn int-IRES, synthetic intron-internal ribosome entry site.

**Figure S5.**
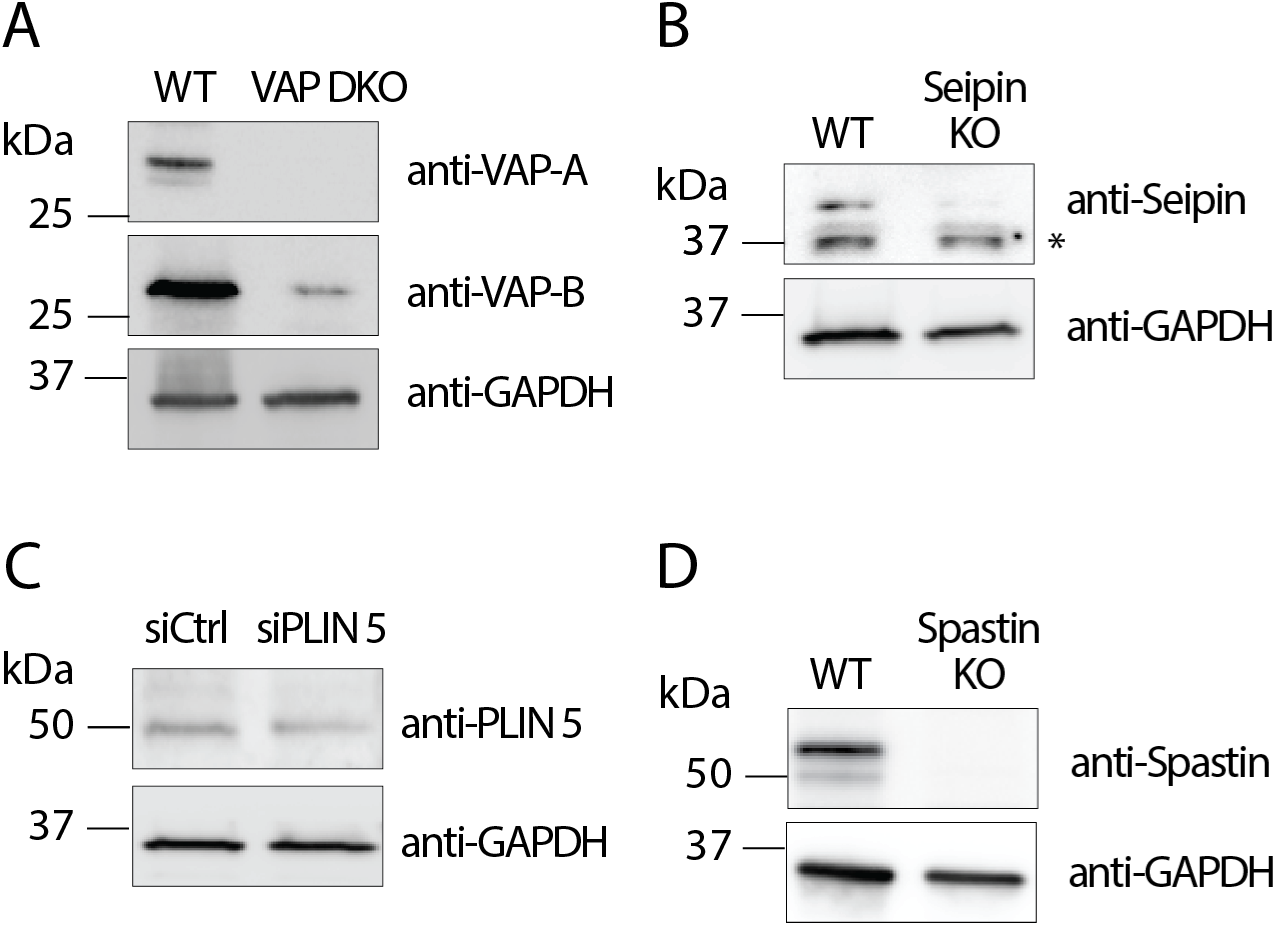
Validation of cell lines via Western blotting. Endogenous protein levels of VAP-A and VAP-B (A), Seipin (B), PLIN5 (C), and Spastin (D) detected by Western blotting. GAPDH protein levels serve as a loading control. * indicates a non-specific band.

**Figure S6.**
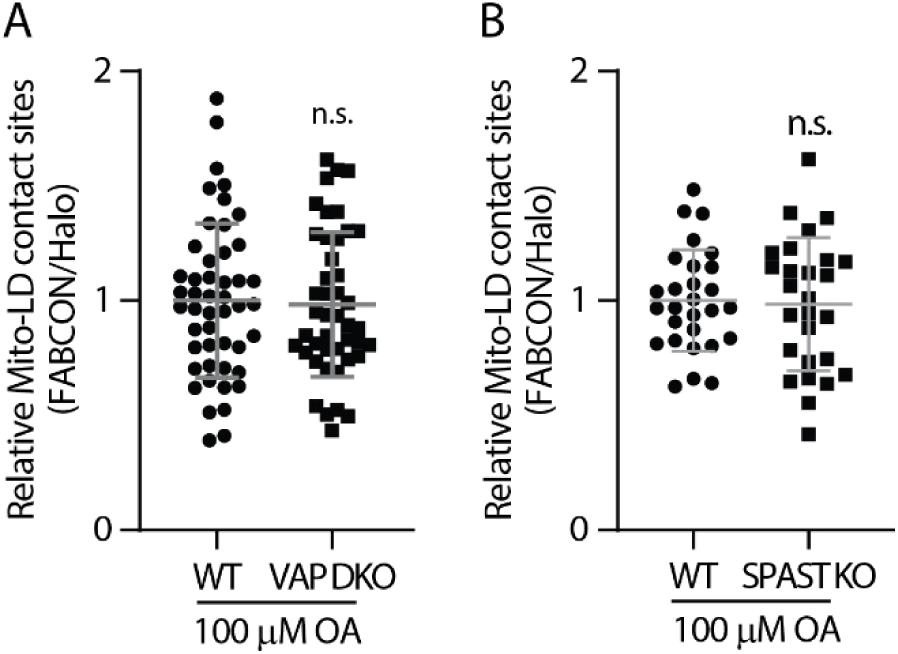
Mito-LD contact sites in VAP DKO and SPAST KO cells. (A) Relative levels of mito-LD contact sites in oleic (OA)-treated WT and VAP DKO HeLa cells. Raw data and mean ± standard deviation are shown (44-48 cells from two independent experiments). (B) Relative levels of mito-LD contact sites in OA-treated WT and SPAST KO U2OS cells. Raw data and mean ± standard deviation are shown (27-28 cells from two independent experiments).

**Table S1.** Organelle targeting motifs.

| Organelle | Targeting motif | Validation | Reference |
| --- | --- | --- | --- |
| Lipid droplet | 6xHp | BODIPY; PLIN2; APEX2 | Shown here |
| ER | Rat Cytochrome b5 TM<br>Sec61β | Calreticulin<br>N/A | (Cho et al., 2020)<br>N/A |
| Mitochondria | Human SYNJ2BP TM | Tomm20 | (Benedetti et al., 2020) |
| Peroxisome | Human full-length PMP34 | Catalase; PMP70 | Shown here |

**Table S2.**
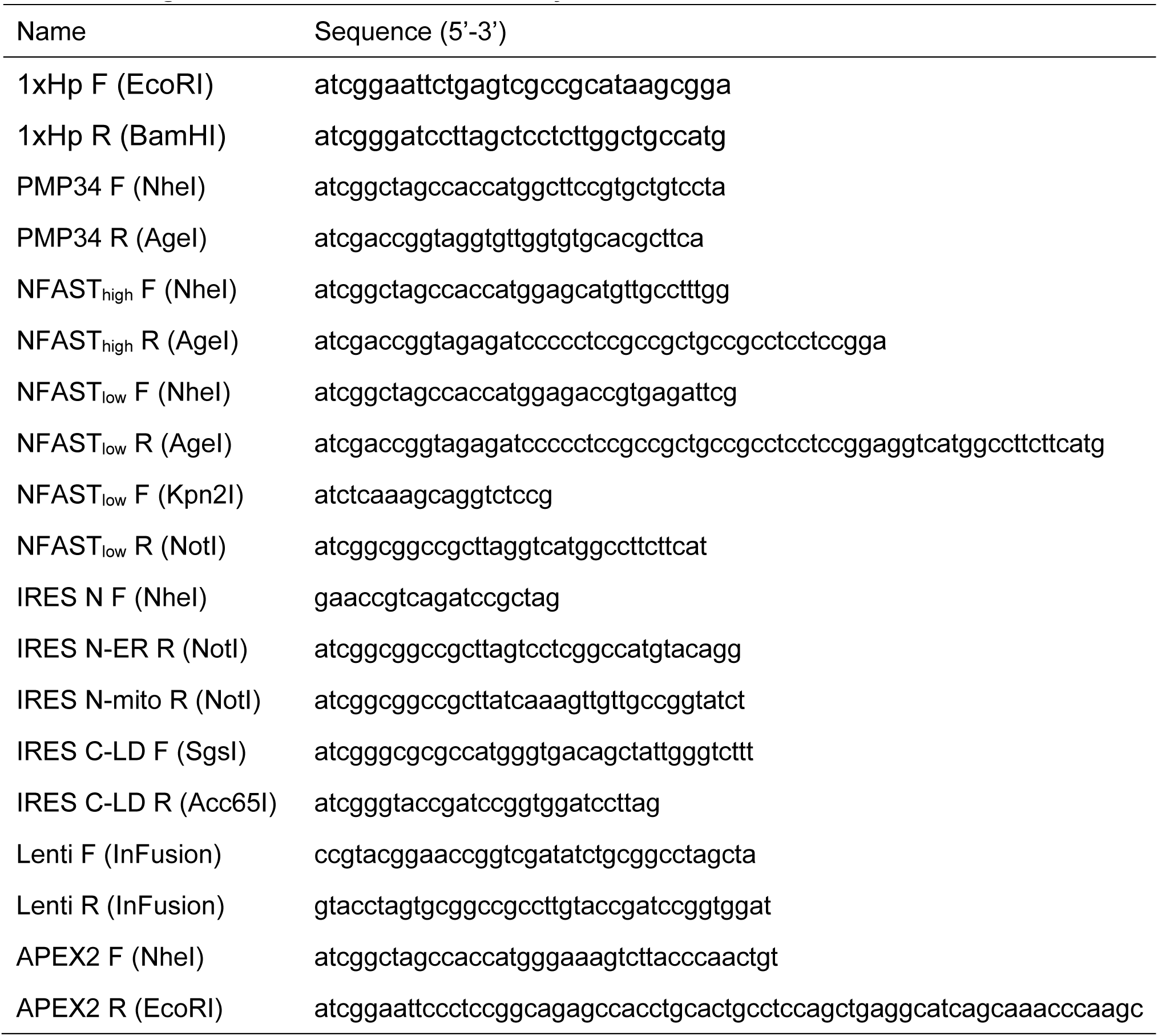
Oligonucleotides used in this study.

